# Nanobody Repertoires for Exposing Vulnerabilities of SARS-CoV-2

**DOI:** 10.1101/2021.04.08.438911

**Authors:** Fred D. Mast, Peter C. Fridy, Natalia E. Ketaren, Junjie Wang, Erica Y. Jacobs, Jean Paul Olivier, Tanmoy Sanyal, Kelly R. Molloy, Fabian Schmidt, Magda Rutkowska, Yiska Weisblum, Lucille M. Rich, Elizabeth R. Vanderwall, Nicolas Dambrauskas, Vladimir Vigdorovich, Sarah Keegan, Jacob B. Jiler, Milana E. Stein, Paul Dominic B. Olinares, Theodora Hatziioannou, D. Noah Sather, Jason S. Debley, David Fenyö, Andrej Sali, Paul D. Bieniasz, John D. Aitchison, Brian T. Chait, Michael P. Rout

## Abstract

Despite the great promise of vaccines, the COVID-19 pandemic is ongoing and future serious outbreaks are highly likely, so that multi-pronged containment strategies will be required for many years. Nanobodies are the smallest naturally occurring single domain antigen binding proteins identified to date, possessing numerous properties advantageous to their production and use. We present a large repertoire of high affinity nanobodies against SARS-CoV-2 Spike protein with excellent kinetic and viral neutralization properties, which can be strongly enhanced with oligomerization. This repertoire samples the epitope landscape of the Spike ectodomain inside and outside the receptor binding domain, recognizing a multitude of distinct epitopes and revealing multiple neutralization targets of pseudoviruses and authentic SARS-CoV-2, including in primary human airway epithelial cells. Combinatorial nanobody mixtures show highly synergistic activities, and are resistant to mutational escape and emerging viral variants of concern. These nanobodies establish an exceptional resource for superior COVID-19 prophylactics and therapeutics.

## INTRODUCTION

SARS-CoV-2, the viral causative agent of COVID-19, has infected over a hundred million people in the last year, killing more than 2.5 million; despite the great promise of vaccines, the resulting pandemic is ongoing. Inequities in vaccine distribution, waning immunity, the biological and behavioral diversity of the human population, and the emergence of viral variants that challenge monoclonal therapies and vaccine efficacy indicate that future outbreaks are highly likely (Diamond et al., 2021; Fraser et al., 2004; Lavine et al., 2021; Wang et al., 2021b; Wang et al., 2021c). Thus, the best we can hope for now is an uneasy truce, in which multi-pronged containment strategies will be required for many years to keep SARS-CoV-2, future variants and novel coronaviruses at bay (McKenna, 2021; Phillips, 2021; Steenhuysen and Kelland, 2021; Weisblum et al., 2020).

Spike (S), the major surface envelope glycoprotein of the SARS-CoV-2 virion, is key for infection as it attaches the virion to its cognate host surface receptor, angiotensin-converting enzyme 2 (ACE2) protein, and triggers fusion between the host and viral membranes, leading to viral entry into the cytoplasm (Walls et al., 2020a; Wrapp et al., 2020b; Zhou et al., 2020). The Spike protein monomer is ∼200☐kDa, extensively glycosylated to help evade immune system surveillance, and exists as a homotrimer on the viral surface. Spike is highly dynamic and is composed of two domains: S1, which contains the host receptor binding domain (RBD); and S2, which undergoes large conformational changes that enable fusion of the viral membrane with that of its host (Hsieh et al., 2020; Letko et al., 2020; Li, 2016; Li et al., 2003; Watanabe et al., 2020). Based on the requirement for attachment to ACE2 for entry, the major target of immunotherapeutics has been the RBD (Barnes et al., 2020; Baum et al., 2020; Finkelstein et al., 2021; Hartenian et al., 2020; Korber et al., 2020; Trigueiro-Louro et al., 2020; Wu et al., 2020).

Major immunotherapeutic strategies to date have focused on immune sera and human monoclonal antibodies; however, these therapies now face the emergence of variants, including RBD point mutants, that have evolved to bypass the most potent neutralizing human antibodies, which are the very basis of immunotherapies (Garcia-Beltran et al., 2021b; Liu et al., 2021b; Starr et al., 2021; Wang et al., 2021c; Weisblum et al., 2020). A specific alternative class of single chain monoclonal antibodies, commonly called nanobodies, can be attractive alternatives to traditional monoclonal antibodies (Muyldermans, 2013). Nanobodies are the smallest single domain antigen binding proteins identified to date, possessing several potential advantages over conventional monoclonal antibodies. Nanobodies are derived from the variable domain (V_H_H) of variant heavy chain-only IgGs (HCAb) found in camelids (e.g. llamas, alpacas, and camels), can bind in modes different from typical antibodies, covering more chemical space and binding with very high affinities (comparable to the very best antibodies) (Jovcevska and Muyldermans, 2020; Muyldermans, 2013). Their small size (~15 kDa) allows them to bind tightly to otherwise inaccessible epitopes that may be (De Genst et al., 2006; Nam et al., 2016) obscured by the glycoprotein coat and so be unavailable to larger antibodies (Laursen et al., 2018; Zare et al., 2021). Nanobodies are highly soluble, stable, lack glycans and are readily cloned and expressed in bacteria (Muyldermans, 2013). They have low immunogenicity (Bannas et al., 2017; Jovcevska and Muyldermans, 2020; Revets et al., 2005) and can be readily modified to be “humanized” (including Fc addition), to change clearance rates, to add cargos such as drugs or fluorophores or to combine and improve characteristics by multimerization (Chanier and Chames, 2019; Duggan, 2018; Vincke et al., 2009). In the case of respiratory viruses like SARS-CoV-2, nanobodies’ flexibility in drug delivery is a critical advantage. Beyond typical administration methods, a major advantage of nanobodies is their potential for direct delivery by nebulization deep into the lungs (Wolfel et al., 2020). This route can provide a high local concentration in the airways and lungs to ensure rapid onset of therapeutic effects, while limiting the potential for unwanted systemic effects (Erreni et al., 2020) as exemplified by clinical trials (Van Heeke et al., 2017; Zare et al., 2021). Nonetheless, many potentially neutralizing nanobodies published to date suffer from the same problem as human monoclonals, in that they also recognize regions of RBD that are subject to escape variation, reducing their potential efficacy (Sun et al., 2021; Wang et al., 2021c).

We refined our methodology to produce large numbers of high affinity nanobodies (Fridy et al., 2014) in order to exploit the available epitope and vulnerability landscape of SARS-CoV-2 Spike protein. The resulting repertoire promises a plethora of synergistically potent and escape resistant therapeutics.

## RESULTS AND DISCUSSION

### Maximizing the Size and Diversity of Anti-SARS-CoV-2 Spike Nanobody Repertoire

We built on our existing pipeline (Fridy et al., 2014), further optimizing each step, explicitly designing it to yield hundreds of high quality, highly diverse nanobody candidates (**Fig. 1A**). Our first improvement involved a pre-screening protocol to select llamas with naturally strong immune responses, as determined by activity against standard animal vaccines (Thompson et al., 2016). After selection, we immunized two llamas with independent Spike subunits, S1 and S2, and used an intensive immunization schedule until we observed excellent responses in both animals, with a strong HCAb component.

**Figure 1.**
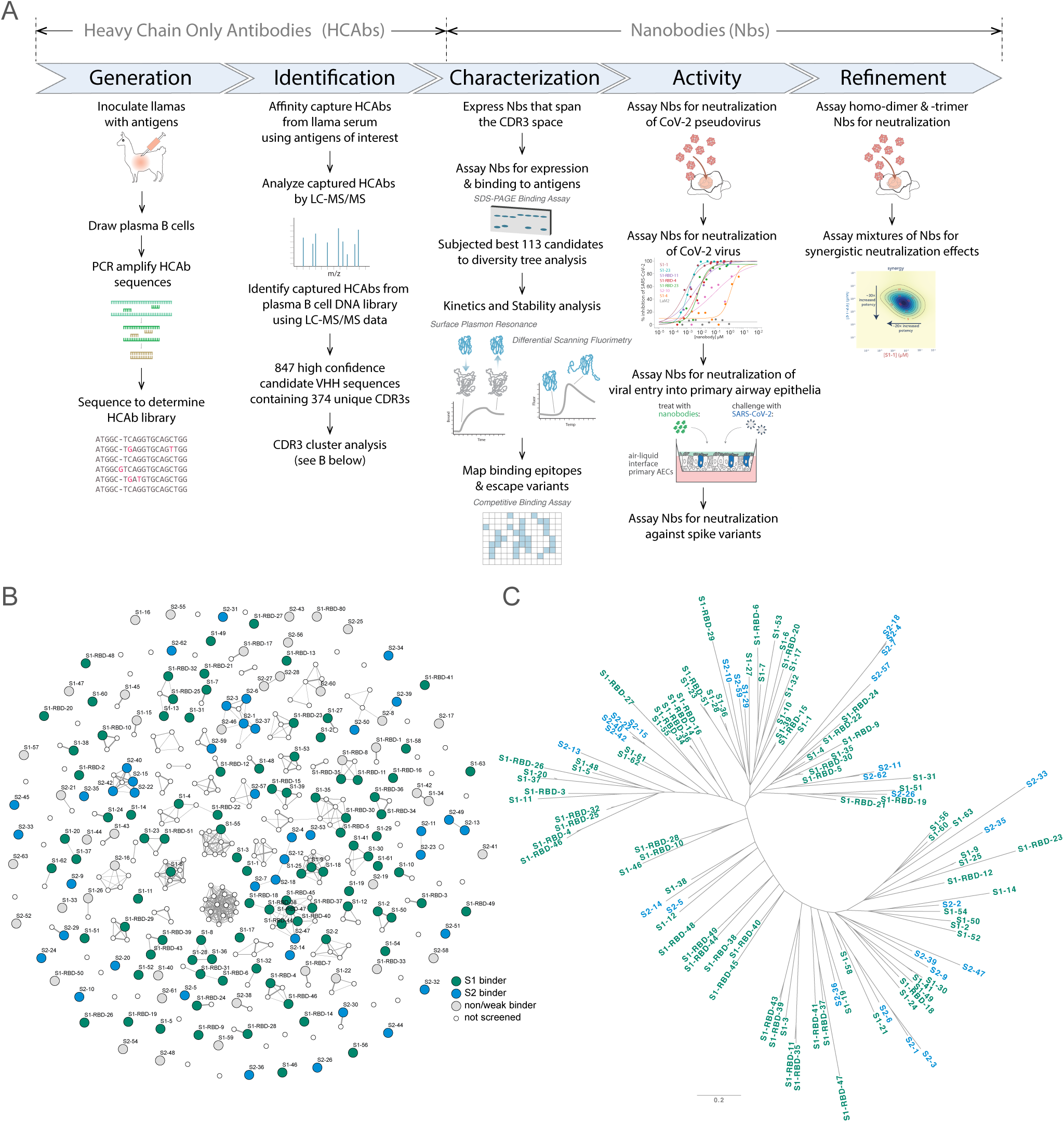
Approach. (A) Schematic of our strategy for generating, identifying and characterizing large, diverse repertoires of nanobodies that bind the Spike protein of CoV-2. The highest quality nanobodies were assayed for their ability to neutralize CoV-2 pseudo-virus, CoV-2 virus and viral entry into primary human airway epithelial cells. We also measured the activities of homodimers/homotrimers and mixtures. (B) A network visualization of 374 high confidence CDR3 sequences identified from Mass Spectrometry workflow. Nodes (CDR3 sequences) were connected by edges defined by a Damerau-Levenshtein distance of no more than 3, forming 183 isolated components. A thicker edge indicates a smaller distance value, i.e. a closer relation. (C) Dendrogram showing sequence relationships between the 113 selected nanobodies, demonstrating that the repertoire generally retains significant diversity in both anti-S1 (green) and anti-S2 (blue) nanobodies, albeit with a few closely related members. Scale, 0.2 substitutions per residue.

To identify V_H_H domains that bind Spike, we affinity purified V_H_H domains from the immunized animals’ sera against Spike S1, S2 or RBD domains, using independent domains in this purification step to maximize epitope accessibility. In parallel, lymphocyte RNA was taken from bone marrow aspirates, and used to amplify V_H_H domain sequences by PCR, which were sequenced to generate an in silico library representative of all V_H_H sequences expressed in the individual animal. The affinity-purified V_H_H fragments were proteolyzed and the resulting peptides analyzed by LC-MS/MS. These data were searched against the V_H_H sequence library to identify and rank candidate nanobody sequences using our Llama-Magic software package (Fridy et al., 2014) with a series of improvements.

To maximize the purity of the serum HCAb sample, we explored different binding conditions to select for the tightest V_H_H binders – a key step not generally available to display panning methods (Fridy et al., 2014). We also used an additional HCAb purification step to deplete VH IgG by incubation with immobilized Protein M, a mycoplasma protein specific for IgG light chain (Grover et al., 2014). To further enrich the V_H_H sample for MS analysis and remove Fc, we perform a digest with IdeS, a protease that cleaves the V_H_H domain from the HCAb with higher specificity than conventionally used papain (von Pawel-Rammingen et al., 2002). Greater peptide coverage for LC-MS was attained by using complementary digestion with trypsin and chymotrypsin (Xiang et al., 2020), augmented by partial SDS-PAGE gel-based separation of different V_H_Hs to reduce the V_H_H complexity and to give more complete peptide coverage and candidate selection. We redesigned PCR primers to maximize coverage of V_H_H sequences for our cDNA libraries. Also, to increase the reliability of the library, singletons were not considered as candidates and priority was given to sequences with high counts. Finally, we updated our Llama-Magic software package (Fridy et al., 2014) to include improved scoring functions, weighting the length, uniqueness and quality of the MS data especially for complementarity-determining regions. This optimized protocol allowed us to identify 374 unique CDR3 sequences (from 847 unique V_H_H candidates).

To maximize sequence diversity and thus the paratope space being explored, we clustered CDR sequences, revealing that many of the candidates form clusters likely to have similar antigen binding behavior. Here, partitioning of the clusters was performed by requiring that CDR3s in distinct clusters differ by a distance of more than three Damerau-Levenshtein edit operations (Bard, 2007) – i.e., each operation being defined by insertion, deletion, or substitution of an amino acid, or transposition of two adjacent amino acids (**Fig. 1B**). This partitioning was found to be effective in that virtually no overlap was observed between those directed against S1 versus S2. The lengths of these CDR3 candidates also varied considerably, ranging from 3 - 22 amino acids in length. The use of two animals further expanded the paratope diversity in that only 4 out of 22 possible clusters from the second animal were observed to be shared with the first animal. In addition, we detected relatively little overlap between our CDR3 clusters and those observed by other groups; for example, just 1 out of 109 S1 specific clusters (Damerau-Levenshtein ≤ 3) were shared by (Xiang et al., 2020) and the present work - indicating that our repertoires are highly orthogonal. This analysis also indicates that we have sampled extended regions of the available paratope space (see also below).

177 high-confidence candidates were selected for expression and screening. Of these, 63 were from S1 affinity purification, 63 from S2, and 51 from RBD, numbered S1-n, S2-n, and S1-RBD-n respectively. These were then expressed with periplasmic secretion in bacteria, and crude periplasmic fractions were bound to the corresponding immobilized spike antigen to assay recombinant expression, specific binding, and degree of binding (**Suppl. Figs. 1,2**). 135 candidates were validated by this screen: 49 against S1, 42 against S2, and 44 against RBD (**Fig. 1B**). To eliminate candidates with the weakest expression and binding affinity, only nanobodies with binding intensity >20% of the observed maximum across all those screened were chosen for follow-up study. This filtering identified the top 113 nanobodies that were purified for further characterization (**Suppl. Tables 1,2**). Note that these selections were designed to provide a strict cutoff in the interests of maximizing the quality of the repertoire selected for thorough characterization, but eliminated many additional bona fide nanobodies that bind to S1 and S2. While a few of these 113 nanobodies were chosen to share similar paratopes, overall, the group retained a high sequence and paratope diversity (**Fig. 1C**).

### High Affinity Nanobodies Across the Entire Spike Ectodomain that are Refractory to Common Spike Escape Mutants

Surface plasmon resonance (SPR) was used to detail the kinetic properties and affinities of the selected nanobodies (**Suppl. Tables 1,2**). All bound with high affinity, with over 60% binding with K_D_s less than 1nM, and two with single digit picomolar affinities (**Fig. 2**). While most S1 binding nanobodies bind RBD (71 nanobodies), 16 nanobodies targeted non-RBD regions of S1 and 26 bind S2 (**Fig. 2**). The disproportionate number of non-RBD S1 and S2 nanobodies reveals the highly antigenic nature of the RBD and highlights likely occlusion of non-RBD S1 regions and S2 due to the glycan shield of SARS-CoV2 Spike (Grant et al., 2020; Watanabe et al., 2020). At the same time, we observed no obvious bias in nanobody affinities for these different domains. While both high on rates and low off rates contributed to these high affinities, kinetic analyses underscore the uniformly fast association rates (k_on_ ≥ 10^+6^) of these nanobodies (likely due to the nanobodies’ small size and proportionally large paratope surface area), with many surpassing the k_on_ rates of high-performing monoclonal antibodies (k_on_=10^+5^) (Tian et al., 2020) (**Fig. 2**), a property that would benefit translation of these nanobodies into rapid therapeutics and diagnostics (Carter, 2006). For those nanobodies with apparently homologous paratopes (**Fig. 2C**), we found no correlation in their kinetic properties (**Suppl. Tables 1,2**), demonstrating that even small paraptope changes can strongly alter behaviors (Fridy et al., 2014).

**Figure 2.**
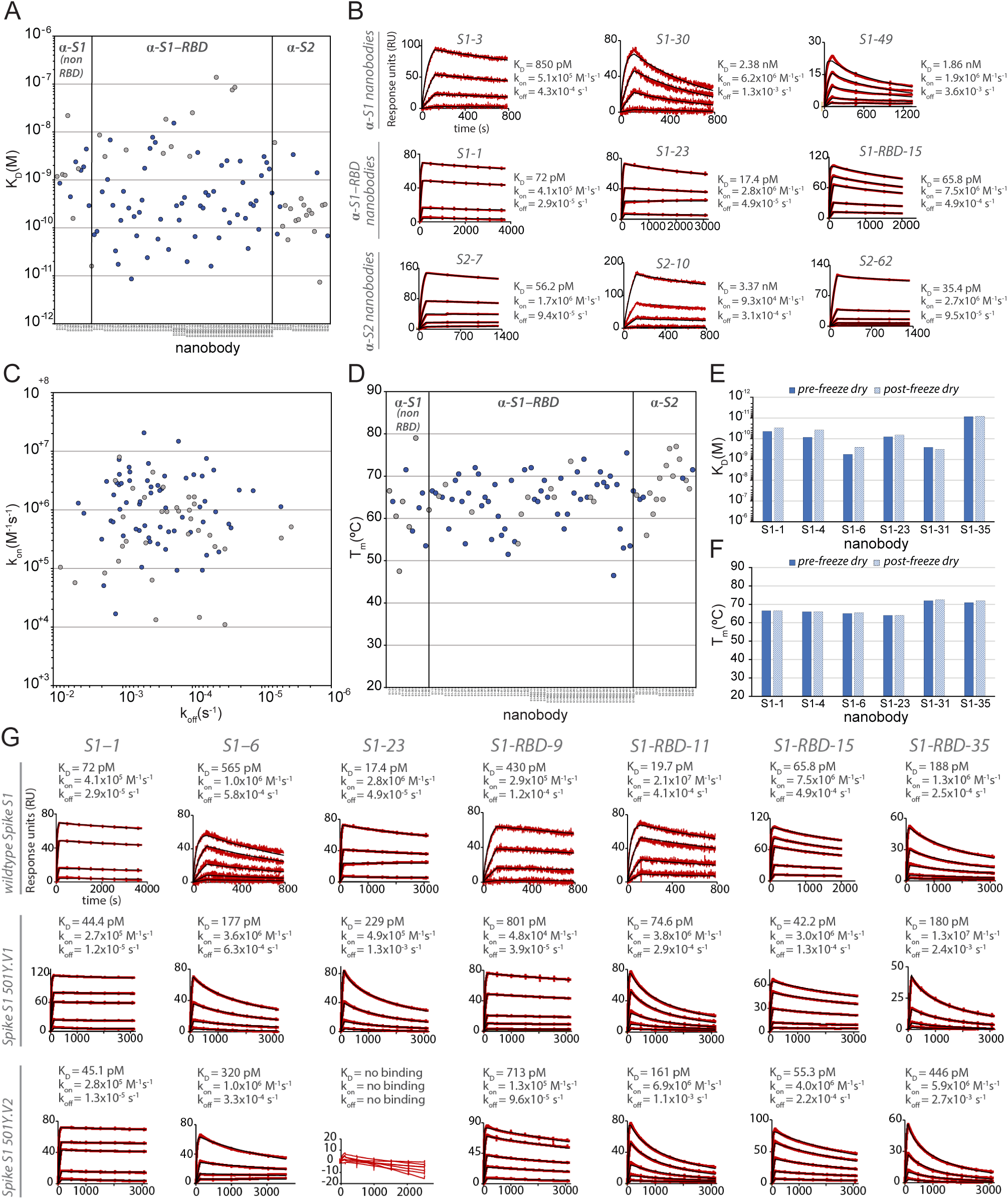
Biophysical characterization of anti-SARS-CoV-2 Spike nanobodies. (A) Each nanobody plotted against their affinity (K_D_) for their antigen separated into three groups based on their binding region on SARS-CoV-2 Spike protein. The data points highlighted in blue correspond to nanobodies that neutralize. The majority of nanobodies have high affinity for their antigen with K_D_s below 1 nm. 10 nanobodies are not included in this plot as they were unable to be analyzed successfully using SPR. (B) SPR sensorgrams for each of the three targets on SARS-CoV-2 Spike protein of our nanobody repertoire, showing three representatives for each binding region. (C) The association rate of each nanobody (k_on_) versus the corresponding dissociation rate (k_off_). The majority of our nanobodies have fast association rates (≥10^+5^ M^-1^s^-1^), with many surpassing the k_on_ of high performing monoclonal antibodies and nanobodies with a k_on_ > M^-1^s^-1^. (D) Each nanobody plotted against their T_m_ as measured by DSF, revealing all but two nanobodies fall within the optimal T_m_ range (between 50°C and 80°C), where the bulk of our nanobodies have a T_m_ ≥ 60°C. No data could be collected for two nanobodies and 10 nanobodies exhibited two dominant peaks in the thermal shift assay and were not included in this plot (a full summary of this data can be seen in **Suppl. Tables 1 and 2**). The K_D_ (E) and T_M_ (F) of six nanobodies was assessed pre- and post-freeze-drying, revealing no significant change in affinity or T_m_ after freeze-drying. (G) SPR sensorgrams comparing the kinetic and affinity analysis of seven nanobodies against wildtype Spike S1 (Wuhan str.), Spike 20I/S1 501Y.V1 (United Kingdom variant) and 20H/Spike S1 501Y.V2 (Sth. African variant).

A worrying recent development has been the emergence of viral variants, with mutations in RBD that minimize or nullify binding of many currently available monoclonal antibodies and nanobodies (Diamond et al., 2021; Garcia-Beltran et al., 2021a; Jangra et al., 2021; Liu et al., 2021a; Sun et al., 2021; Wang et al., 2021c; Weisblum et al., 2020). Indeed in one study, the efficacy of 14 out of the 17 most potent monoclonal antibodies tested was compromised by such common RBD mutants (Wang et al., 2021c). Here, nanobodies show great potential to be particularly resistant to these variants (Sun et al., 2021). RBD mutants represent a significant class of escape variants (Garcia-Beltran et al., 2021b; Greaney et al., 2021), and so two strategies were employed to ensure the generation of numerous nanobodies whose binding (and virus neutralizing activities) are also resistant to emerging variants. First, we isolated a large diversity of high quality anti-RBD nanobodies to maximize the probability of identifying ones that are refractory to escape. Second, we targeted non-RBD regions of Spike (see below) (Elshabrawy et al., 2012; Greaney et al., 2021). To test the first strategy, we sampled RBD binding nanobodies covering non-overlapping epitopes on RBD (see below) and examined their binding to SARS-CoV2 variants B.1.1.7 / 20I/501Y.V1 (United Kingdom) and B.1.351 / 20H/501Y.V2 (South Africa), currently among the most prevalent and concerning among those spreading in the population (Ho et al., 2021; Wang et al., 2021b) (**Fig. 2**). Of the seven nanobodies tested, six of these (S1-1, S1-6, S1-RBD-9, S1-RBD-11, S1-RBD-15 and S1-RBD-35) retained their very strong binding to both variants, with only a modest reduction in affinity for S1-RBD-11 binding to variant B.1.351 (20 pM to 161 pM). For the seventh nanobody, S1-23, binding to variant B.1.1.7 / 20I/501Y.V1 was only reduced from a K_D_ of 17 pM to a still-respectable 230 pM, however, its binding to variant B.1.351 / 20H/501Y.V2 was abolished (**Fig. 2**). As expected (Magnus, 2013; Steckbeck et al., 2005; VanCott et al., 1994), it is the off rates that are most affected by these variants. Nevertheless, based on epitope mapping (below) and our identification of nanobodies that recognize epitopes not altered in the emerging variant strains, we expect that a high percentage of our nanobodies will remain resistant to these escape mutants, making our collection a powerful resource for potential prophylactics and therapeutics.

### The Nanobody Repertoire Has Favorable Stability Properties

A key consideration for possible biological therapeutics and diagnostics for SARS-CoV-2 is their stability under potentially denaturing conditions (McConnell et al., 2014). To address this, differential scanning fluorimetry (DSF) experiments were performed to determine the thermal stability (*T*_m_) of each of our nanobodies. These studies revealed a thermal stability range between 50°C and 80°C, similar to published results of other properly folded nanobodies and indicative of the high stability generally associated with nanobodies (Muyldermans, 2013). In contrast to many conventional antibodies, nanobodies are also reported to remain fully active upon reconstitution after lyophilization, particularly in buffers lacking cryoprotectants (Schoof et al., 2020b; Xiang et al., 2020). A representative sample from our repertoire was thus freeze-dried without cryoprotectants, reconstituted, then analyzed via SPR and DSF to determine whether their properties were compromised due to lyophilization. The results revealed no significant effect on stability, kinetics and affinity (**Fig. 2**). Taken together, these data suggest that our nanobodies, like those published in other contexts (Schoof et al., 2020a; Xiang et al., 2020) are highly robust and able to withstand various temperatures and storage conditions without affecting their stability and binding. These are essential requirements for downstream applications (e.g. use in a nebulizer) and ease of storage - important considerations if these are to be used for mass distribution to populations across the globe, including in resource-poor settings (Peeling and McNerney, 2014).

### Nanobodies Explore the Available Epitope Landscape of the Spike Ectodomain

We applied a multifaceted approach to physically distinguish nanobodies that target common regions on the surface of the RBD. Using an eight-channel bio-layer interferometer we tested subsets of our RBD-specific nanobodies for pair-wise competitive binding to the RBD (**Fig. 3**). Label-free binding of antibodies to antigens measured in a “dip-and-read” mode provides a real-time analysis of affinity and the kinetics of the competitive binding of nanobody pairs and can distinguish between those that bind to similar or overlapping epitopes versus distinct, non-overlapping epitopes (Estep et al., 2013). 44 anti-RBD nanobodies were screened in pairwise combinations. The response values were used to assist the discovery of nanobody groups that most likely bind non-overlapping epitopes, by ensuring that the least response of pair-wise nanobodies within the group is maximized. Nine representative nanobodies from this group were used as a foundation, selecting two or more representative nanobodies from each group to bin the remaining RBD nanobodies in our collection. Overlapping pairs from the foundation group and the remaining RBD binders were used to measure if a nanobody pair behaved similarly against other nanobodies measured in the dataset (**Fig. 3A**), to comprehensively map nanobody competition and epitope bins (**Fig. 3B**). Pearson correlation coefficients were derived based on their binding characteristics and the data were used to hierarchically cluster and group all RBD binders into bins. This approach revealed three large, mostly non-overlapping bins. However, each bin contained smaller, better-correlated clusters of nanobodies, reflected by the dendrogram, indicating the presence of numerous distinct sub-epitope bins present within each larger bin, i.e. discrete epitopes that partially overlap with other discrete epitopes in the same bin. We calculated the gap statistic (Tibshirani et al., 2001), which optimizes the estimation of the optimal cluster number, discerning at least 15 epitope bins (**Fig. 3A**).

**Figure 3.**
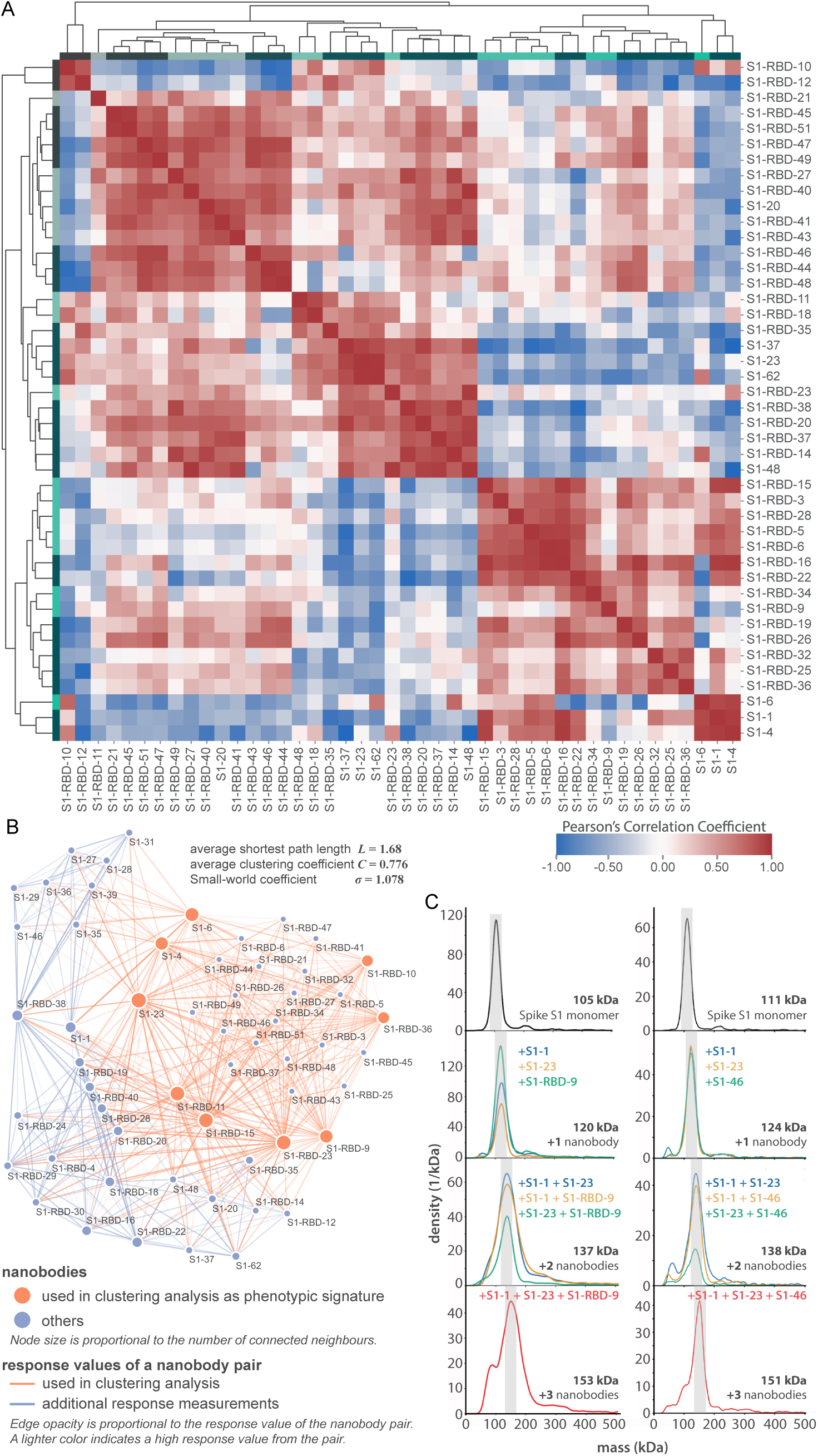
Epitope characterization of nanobodies against the S1-RBD of SARS-CoV-2 Spike. (A) Major epitope bins are revealed by a clustered heat map of Pearson’s Correlation Coefficients computed from the response values of nanobodies binding to the Spike RBD in pairwise cross-competition assays on a biolayer interferometer. Correlated values (red) indicate that the two nanobodies respond similarly when measured against a panel of nine RBD nanobodies that bind to distinct regions of the RBD. A strong correlation score indicates binding to a similar/overlapping region on the RBD. Anti-correlated values (blue) indicate that a nanobody pair responds divergently when measured against nanobodies in the representative panel, and indicate binding to distinct or non-overlapping regions on the RBD. Hierarchical clustering analysis reveals 15 separate and partially overlapping epitope bins as visualized by alternating dark and light teal bars linked together by a dendrogram. (B) A network visualization of anti-S1-RBD nanobodies. Each node is a nanobody and each edge is a response value measured by biolayer interferometry from pairwise cross-competition assays. Orange nodes represent nine nanobodies used as a representative panel for clustering analysis in A. Blue nodes represent the other anti-S1-RBD nanobodies present in the dataset. The average shortest distance between any nanobody pair in the dataset of 1.68. An average clustering coefficient of 0.776 suggests that the measurements are well-distributed across the dataset. The small world coefficient of 1.078 indicates that the network is more connected than to be expected from random, but the average path length is what you would expect from a random network, together indicating that the relationship between nanobody pairs not actually measured can be inferred from the similar/neighboring nanobodies. (C) Mass photometry (MP) analysis of Spike S1 monomer incubated with different anti-Spike S1 nanobodies. Two examples of an increase in mass as Spike S1 monomer (black line) is incubated with one to three nanobodies. The accumulation in mass upon addition of each different nanobody on Spike S1 monomer is due to each nanobody binding to non-overlapping space on Spike S1, an observation consistent with Octet binning data. As a control, using MP, each individual nanobody was shown to bind Spike S1 monomer on its own (data not shown).

The binning data from pairwise combinations suggest numerous epitope bins, and thus it is reasonable to hypothesize that more than two nanobodies can bind to the S1-RBD at the same time. To test this hypothesis we used mass photometry (MP), which can accurately measure multiple binding events to a single antigen, allowing us to determine which nanobodies share epitope space on Spike S1 monomer through detection of additive mass accumulation of a nanobody (or nanobodies) on Spike S1 depending on whether or not nanobodies share epitope space on Spike S1. Several representative nanobodies that sample across the epitope space of our nanobody repertoire were chosen for MP studies based on the epitope binning data. These data confirmed the separation of our major epitope bins, and furthermore demonstrated that we can bind at least three different nanobodies simultaneously to the RBD of S1, an important consideration for the design of complementary and synergistic nanobody cocktails and multimers (**Fig. 3C**).

### Anti-RBD Nanobodies are Highly Effective Neutralizing Agents

We used a SARS-CoV-2 pseudovirus neutralization assay to screen and characterize our nanobody repertoire for antiviral activities (**Fig. 4**). The lentiviral-based, single round infection assay robustly measures the neutralization potential of a candidate nanobody and is a validated surrogate to replication competent SARS-CoV-2 (Riepler et al., 2020; Schmidt et al., 2020). Overall, ~40% of our monomeric nanobody repertoire neutralized with IC50s <100 nM, while 26% showed neutralization with IC50s <50 nM and 16 potent neutralization at 20 nM or lower (**Fig. 4A**). Because measured IC50s are dependent on assay conditions and so cannot be readily compared across laboratories (Cheng and Prusoff, 1973), we included, as benchmarks, other published nanobodies (Wrapp et al., 2020a; Xiang et al., 2020) (**Fig. 4G**). These four selected nanobodies were cloned and produced in-house and span the range of neutralization observed within our repertoire from potent (<20 nM) to relatively weak (between 1-10 µM). The most potent neutralizing nanobodies mapped to the RBD; neutralizing activity mapped to each of the major epitope bins of the RBD and were of similar efficacy to benchmarks.

**Figure 4.**
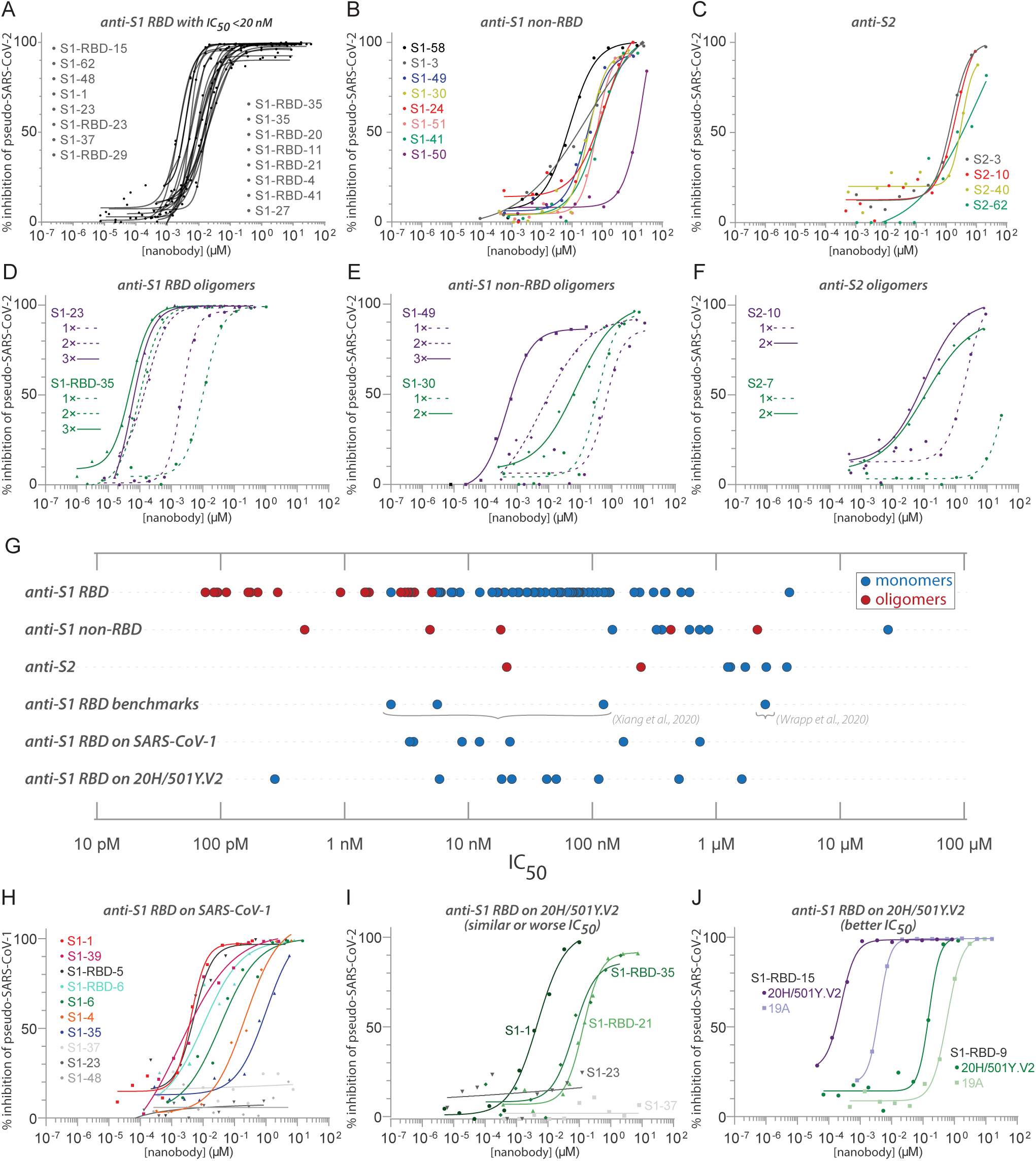
Diverse and potent nanobody-based neutralization of SARS-CoV-2. Nanobodies targeting the S1-RBD, S1 non-RBD, and S2 portions of Spike effectively neutralize lenti-virus pseudotyped with various SARS-CoV Spikes and their variants from infecting ACE2 expressing HEK293T cells. (A) Of the 113 nanobodies, monomers that neutralize SARS-CoV-2 pseudovirus with IC50 values 20nM and lower are displayed. (B) Representative nanobodies targeting the non-RBD portions of S1 and (C) the S2 domain of SARS-CoV-2 neutralize SARS-CoV-2 pseudovirus. (D-F) Oligomerization of RBD, S1 non-RBD and S2 nanobodies significantly increases neutralization potency. (G) Summary scatter plot of all nanobody IC50s across the major domains of SARS-CoV-2 Spike and where tested, across SARS-CoV-2 variant 20H/501Y.V2 and SARS-CoV-1. Representative published nanobodies were also tested in our neutralization assays and show similar potency towards SARS-CoV-2 pseudovirus. (H) Representative SARS-CoV-2 RBD targeting nanobodies cross-neutralize SARS-CoV-1 pseudotyped lentivirus and (I-J) the 20H/501Y.V2 SARS-CoV-2 variant (B.1.351) with L18F, D80A, K417N, E484K, and N501Y amino acid substitutions in Spike. 19A (I-J) is the initial SARS-CoV-2 clade that includes the prototypical Wuhan-Hu-1 Spike used as “wild-type” in these pseudovirus assays. In all cases, n>/=2 biological replicates of each nanobody monomer/oligomer with a representative biological replicate with n=4 technical replicates per dilution displayed.

### Nanobody-based Neutralization Beyond the RBD

Notably, nanobodies mapping outside of the RBD on S1 (anti-S1, non RBD) and mapping to S2 also neutralized the pseudovirus in this assay, albeit with somewhat higher IC50s (≳ 150nM for anti-S1, non RBD, e.g. S1-58; and ≳1µM for anti-S2, e.g. S2-3) (**Fig. 4B,C**). Thi is the first evidence of nanobody neutralization activity mapping outside of the RBD. As nanobodies are monomeric, the mechanism of this neutralization does not involve viral aggregation and likely reflects disruption of the viral binding or spike driven fusion of viral and cellular membranes of fusion with target cell membranes. Such nanobodies, especially directed against relatively invariant regions of coronavirus Spike proteins, may have broad spectrum activities.

Further optimization, including testing cocktails and oligomers, is impractical for all 113 nanobodies, due to combinatorial scaling issues - for example, simple binary mixtures of all would number >10^4^ combinations. We therefore carefully selected subsets of nanobodies for such testing based on their favorable properties and epitope coverage.

### Oligomerization Strongly Enhances the Affinity and Neutralization Activity of Nanobodies

A distinct advantage of nanobodies is the facility by which oligomers can be generated and expressed to improve their affinities and avidities. Oligomerization of most nanobodies tested improved their IC50s and measured affinities. Monomeric S1-RBD-35 was converted to dimers and trimers, improving its neutralization activity from IC50s of ~12nM to ~160pM and ~75pM, respectively. Similar results were found with S1-23, improving neutralization from ~7nM to ~170pM and ~90pM, respectively (**Fig. 4D).** The simple prediction that multimerization improves efficiency was not always the case (data not shown). Dimerization of the anti-S1 non RBD nanobody S1-49 improved IC50s from ~1µM to ~5nM, trimerization improved its activity an additional ~5-fold. Multimerization of some nanobodies directed against regions outside of the RBD on both S1 and S2 led to nanomolar range IC50s (**Fig. 4E,F**). This includes S2-7, for which dimerization converted a nanobody that we considered to be a non-neutralizer to one having a respectable neutralizing activity (IC50 ~ 250nM) (**Fig. 4F**). These results show that multimerization can have a dramatic effect on activity, although currently this must be determined empirically.

### Nanobodies Neutralize SARS-CoV-1 and SARS-CoV-2 variants

Both SARS-CoV-1 and SARS-CoV-2 share the same host receptor, ACE2, and the RBDs of the viruses share ~74% identity. As a result, some antibodies and nanobodies have been shown to be cross-neutralizing (Liu et al., 2020a; Wrapp et al., 2020a). We therefore tested the ability of our nanobodies to neutralize SARS-CoV-1 in the pseudovirus assay. Of the nanobodies tested in this assay, numerous (8 of 23 tested) of our anti-RBD monomer nanobodies also displayed excellent neutralizing activities against SARS-CoV-1 Spike pseudotyped virus (**Fig. 4H**). While some nanobodies such as S1-35 and S1-6 showed reduced activity against SARS-CoV-1 pseudotypes compared to those pseudotyped with SARS-CoV-2 Spike, S1-1, −39 and S-1-RBD-5 −6 had similar IC50s against both pseudotypes. Notably, S1-23, −37 and −48 showed no activity against SARS-CoV-1 Spike pseudotypes, all of these being highly correlated with one another in the epitope binning analysis, therefore likely targeting proximal epitopes on Spike (**Fig. 3A**). Beyond nanobodies that bind to the RBD, 2 of 8 nanobodies that bind to non-RBD regions of S1 and S2 also neutralized SARS-CoV-1 Spike pseudotypes (**Suppl. Table 5**).

Certain mutations appearing in ‘variants of concern’ (VOC) have been associated with rapidly increasing case numbers in certain locales, have been demonstrated to reduce the neutralization potency of some monoclonal antibodies and polyclonal plasma, increase the frequency of serious illness, and are spreading rapidly (19A; Wuhan-Hu-1) virus (Wang et al., 2021c; Wibmer et al., 2021). We therefore tested a subset of our better neutralizing nanobodies against a pseudovirus carrying the Spike protein of the B.1.351 / 20H/501Y.V2 variant, that includes the three amino acid substitutions at positions K417N/T, E484K and N501Y (Stamatatos et al., 2021; Tegally et al., 2021). While nanobodies S1-23 and S1-37 failed to neutralize the pseudovirus variant, nanobody S1-1 was equally efficacious against the variant as against the pseudovirus carrying wild-type Spike (**Fig. 4I**). Both S1-RBD-21 and −35 also remained effective neutralizers of the 20H/501Y.V2 variant Spike pseudotypes, albeit with ~10-fold reduction in their IC50s compared to the wild-type Spike (**Fig 4I)**. Remarkably though, two nanobodies, S1-RBD-9 and −15 showed *increased* neutralization activity against the 20H/502Y.V2 Spike pseudoviruses, with the activity of S1-RBD-15 increasing ~10-fold (**Fig. 4J**). These results are also in accord with our Spike variant SPR studies which showed that S1-1, S1-RBD-9, −15, and −35 retained very strong binding to the SA variant (B.1.351 / 20H/501Y.V2, K417N E484K N501Y), whereas binding of S1-23 was completely abolished (**Fig. 2G**). Overall, these data suggest that comprehensive mining of our repertoire and multimerization can lead to nanobody-based therapies that remain fully effective against common and potentially yet-to-emerge variants of SARS-CoV-2 and with broad spectrum coronavirus inhibition activities.

### Nanobodies Effectively Neutralize SARS-CoV-2 Infection in Human Primary Airway Epithelium

Nanobody and antibody neutralizations have been reported to yield similar results when performed with pseudovirus versus authentic virus (Schmidt et al., 2020; Schoof et al., 2020a; Xiang et al., 2020). However, discrepancies have also been reported, particularly for antibodies targeting regions outside the RBD (Chi et al., 2020; Huo et al., 2020). We therefore selected a panel of exemplar nanobodies to test for neutralization with authentic SARS-CoV-2. All nanobodies that neutralized pseudovirus also showed potent neutralization by plaque and focus reduction assays and correlated well with our pseudovirus assays (**Fig. 5A**).

**Figure 5.**
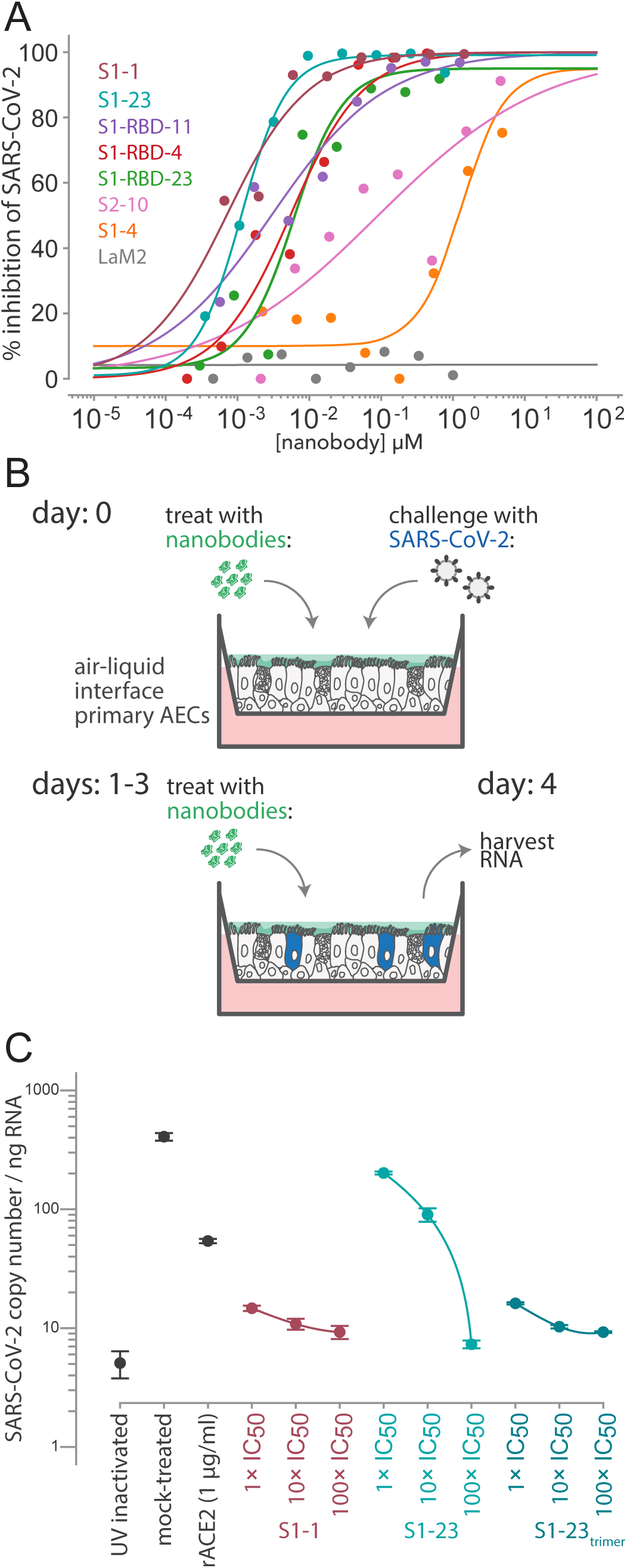
Authentic SARS-CoV-2 neutralization by anti-Spike nanobodies. (A) Neutralization curves are plotted from the results of a focus-forming reduction neutralization assay with the indicated nanobodies. Serial dilutions of each nanobody were incubated with SARS-CoV-2 (MOI 0.5) for 60 min and then overlaid on a monolayer of Vero E6 cells and incubated for 24 h. LaM2, an anti-mCherry nanobody (Fridy et al., 2014) was used as a non-neutralizing control. After 24 h, cells were collected and stained with anti-Spike antibodies and the ratio of infected to uninfected cells was quantified by flow cytometry. (B) A schematic of an air-liquid interface (ALI) culture of primary human airway epithelial cells (AECs) as a model for SARS-CoV-2 infection. Cells were incubated with nanobodies and then challenged with SARS-CoV-2 (MOI 0.5). After daily treatment with nanobodies for three more days, the cultures are harvested to isolate RNA and quantify the extent of infection. (C) AECs were infected with the indicated concentrations of anti-SARS-CoV Spike nanobodies. The infected cultures were maintained for five days with a daily 1 h incubation of nanobodies before being harvested for RNA isolation and determination of the SARS-CoV-2 copy number by qPCR. SARS-CoV-2 copy number was normalized to total RNA measured by spectrophotometry. Mock-treated samples exposed to infectious and UV-inactivated SARS-CoV-2 virions served as positive and negative controls. Recombinant soluble angiotensin converting enzyme 2 (rACE2) was used as a positive treatment control. The indicated nanobodies were used at 1, 10, and 100x their IC50 values determined in pseudovirus neutralization assays.

We also tested a subset of our nanobodies in a human *ex vivo* model system that represents the initial site of SARS-CoV-2 infection and would reflect the ability of our nanobodies to block SARS-CoV-2 infection and spread (**Fig. 5B**). Air-liquid interface (ALI) cultures of primary airway epithelium mimic the lung environment as pseudostratified, ciliated, and mucous secreting cells that express ACE2 (Murphy et al., 2020). SARS-CoV-2 readily infects and replicates in this system (Barrow et al., 2021). We treated the air-exposed apical surface of the culture with serial dilutions of S1-1 and S1-23 and then challenged them with SARS-CoV-2 at an MOI of 0.5. To simulate a treatment regimen, we further treated the ALI cultures with nanobodies at 24 h intervals for an additional 3 days before harvesting the cells, extracting RNA, and measuring SARS-CoV-2 levels by qPCR (**Fig. 5B**). S1-1 potently neutralized SARS-CoV-2 at each concentration tested while S1-23 inhibited SARS-CoV-2 in a dose dependent manner (**Fig. 5C**). The efficacy of the S1-23 nanobody was strongly enhanced when provided to cells as a trimer, potently inhibiting viral replication at levels comparable to those observed with monoclonal antibodies cloned from convalescent serum (**Fig. 5C**) (Robbiani et al., 2020; Seydoux et al., 2020; Stamatatos et al., 2021; Suthar et al., 2020) and these data highlight the potential for nanobodies to function in a therapeutic capacity. As an additional comparator and as a control, we determined the inhibition of replication upon addition of recombinant competitor, ACE2. Nanobodies inhibited at lower doses than recombinant ACE2, reflective of our measured low K_D_ of nanobody interactions with Spike (< 1nM) compared to a reported K_D_ of 14.7 nM or greater for ACE2 with Spike (Cao et al., 2020; Chan et al., 2020; Huang and Chai, 2020; Liu et al., 2020b; Rogers et al., 2020; Shang et al., 2020).

### Synergistic Activity with Nanobody Combinations

Drugs are often combined to improve single drug efficacies and to reduce efficacious drug concentrations. Synergy occurs when the combination of drugs has a greater effect than the sum of the individual effects of each drug. A major advantage of a large repertoire of nanobodies that bind to different epitopes on Spike is the potential for cooperative activity among nanobody pairs (or higher order combinations) leading to synergistic effects. Because our repertoire provides for thousands of pairwise combinations and this scales exponentially when considering cocktails with three or more nanobodies, we tested pairs of nanobodies for synergistic activities. To select nanobody pairs, we took advantage of our epitope mapping, structural data and biophysical performance of nanobodies. Using an automated platform we titrated pairwise combinations of nanobodies in a 2D dilution format and measured their IC50s in the pseudovirus assay. IC50s were modeled using the synergy framework, multi-dimensional synergy of combinations (MuSyC), which models a two-dimensional (2D) Hill equation and extends it to a 2D surface plot. Synergy is evidenced by the parameters of the modeled Hill function. We focused on combinations of nanobodies with S1-23, which itself has a potent IC50 (**Fig. 4A**; **Suppl. Table 1**). Combinations of S1-27 and S1-23 showed simple additive effects (**Fig. 6A**). These nanobodies belong to the same epitope bin (Fig. 2B); their additive effect is as expected for two nanobodies accessing the same site on S1-RBD, but effectively doubling the concentration of a single nanobody.

**Figure 6.**
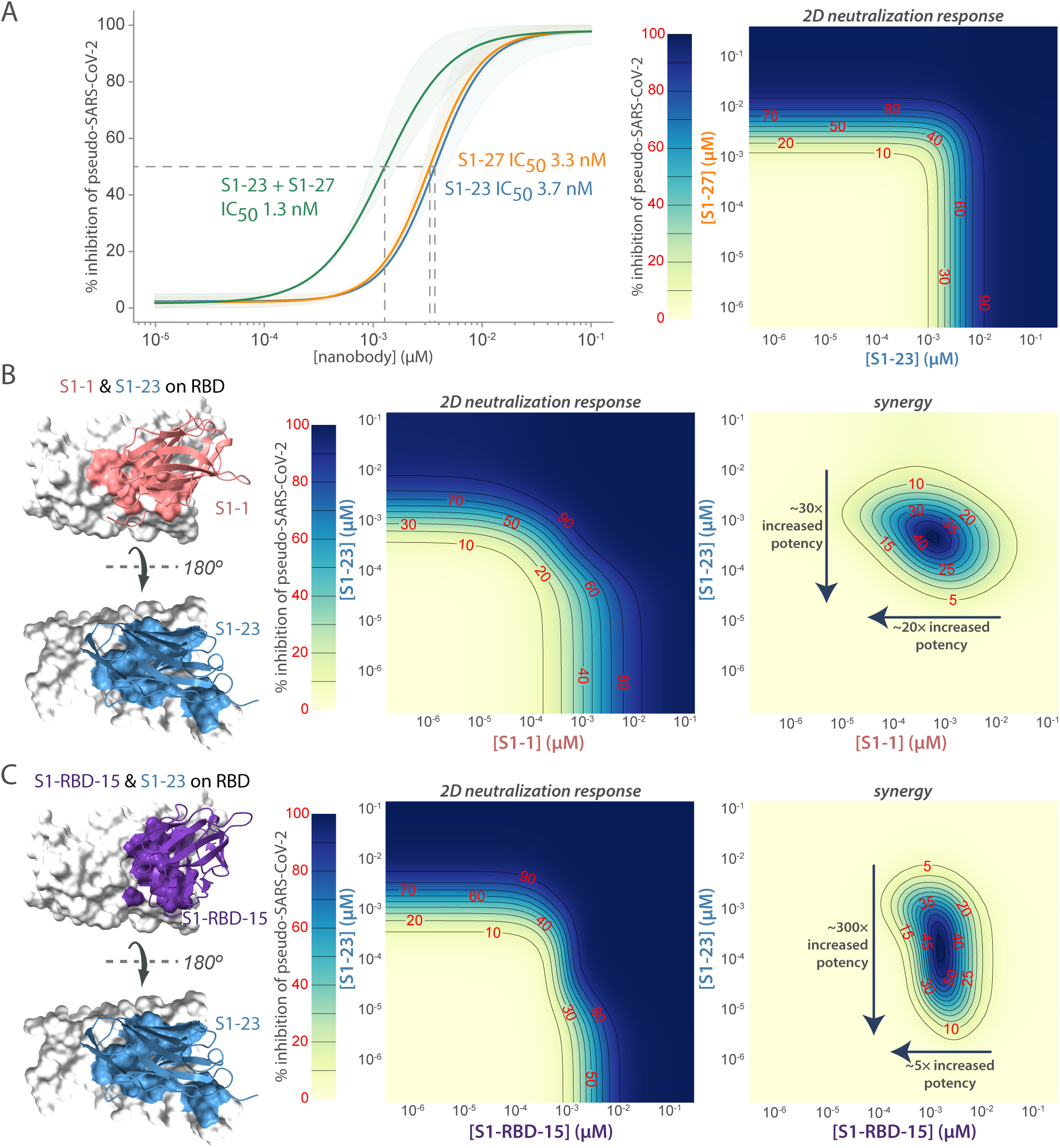
Synergistic neutralization of Spike with nanobody cocktails. (A) An example of additive effects between two anti-SARS-CoV2 Spike nanobodies. S1-23 and S1-27 were prepared in a two-dimensional serial dilution matrix and then incubated with SARS-CoV-2 pseudovirus for 1 h before adding the mixture to cells. After 56 h, the expression of luciferase in each well was measured by addition of Steady-Glo reagent and read out on a spectrophotometer. Neutralization curves and the calculated IC50 of each nanobody alone, or in a 1:1 combination was determined. The right panel shows a heat map of pseudovirus neutralization by a two-dimensional serial dilution of combinations of S1-23 and S1-27. Lines and red numbers demarcate the % inhibition, that is, inhibitory concentration where X% of the virus is neutralized, e.g. IC50. Dark blue regions are concentrations that potently neutralize the pseudovirus, as per the heat map legend. (B) The left panel shows a model of S1-1 and S1-23 neutralizing nanobodies binding to distinct epitopes of the RBD. The middle panel shows the heatmap of pseudovirus neutralization observed by a two-dimensional serial dilution of combinations of S1-1 and S1-23. The right panel shows a heat map with the difference between the observed neutralization and that expected in a null model of only additive effects. The lines and red numbers demarcate regions in the heat map where the observed neutralization is greater than additive by the indicated percentages (red numbers). Overall, S1-1 enhances the effect of S1-23 ~30-fold, whereas S1-23 enhances the effect of S1-1 ~20 fold. (C) As in B, but comparing S1-RBD-15 with S1-23. The left panel shows a model of S1-RBD-15 and S1-23 neutralizing nanobodies binding to distinct epitopes of the RBD. The middle panel shows a heatmap of pseudovirus neutralization observed for 2D serial dilution of S1-RBD-15 and S1-23. The right panel shows the synergy observed between S1-RBD-15 and S1-23. Overall, S1-RBD-15 enhances the effect of S1-1 ~300 fold, whereas S1-23 enhances the effect of S1-RBD-15 ~5 fold.

To move beyond additive effects, we need to consider combinations that bind to different epitopes. Indeed, synergistic effects were observed between S1-23 and S1-1 or S1-RBD-15, or S2-40 and S1-23 (**Fig. 6B,C**). The most dramatic synergistic effect was observed with S1-RBD-15 which improves the potency of S1-23 by ~300-fold. In order to more precisely map epitopes and determine binding modes on the RBD domain, we integrated information from cross-linking mass spectrometry, shape complementarity of nanobody and antigen, locations of escape mutations, and ability (or lack of ability) of different nanobodies to bind RBD simultaneously, to structurally model three RBD-nanobody complexes (namely S1-1, S1-23 and S1-RBD-15). First, we cross-linked nanobody-RBD complexes using disuccinimidyl suberate (DSS) and identified intermolecular crosslinks by MS. Next, we developed comparative structural models of the three aforementioned nanobodies. The ensemble of docked nanobody models satisfied all 27 crosslinks used in the structural modeling (**Suppl. Table 6**). Interestingly, nanobodies with synergistic neutralizing activity show nearby, but non-overlapping epitopes on the RBD of Spike (**Fig. 6B,C**), and can provide a roadmap to the rational production of multimeric, even higher affinity reagents capable of neutralization at low doses while minimizing susceptibility to escape mutations.

### Escape-Resistant Nanobody Cocktails

With the emerging variants of concern, our goal is to develop nanobody multimers and cocktails that are maximally refractory to escape by such variants. To do so, we used a previously employed method that drives the selection of antibody resistant populations of rVSV/SARS-CoV2 chimeric virus harboring variants of Spike and measured the ability of the chimeric virus to escape nanobody-mediated neutralization (Weisblum et al., 2020). This approach simultaneously maps the escape potential of Spike and the epitopes responsible for neutralization by nanobody binding, with the goal of discovering Spike variants that resist the neutralizing activity of individual nanobodies. Based on this information, we could then predict pairs of nanobodies whose escape mutants do not map to the same region of Spike, the combination of which would thus likely prevent escape. Specifically, we prepared large and diversified populations (10^6^ infectious units) of a recombinant VSV derivative (rVSV/SARS-CoV-2/GFP wt_2E1_) that encode SARS-CoV2 Spike protein in place of VSV-G, and recapitulates the neutralization properties of authentic SARS-CoV-2 (Schmidt et al., 2020). The rVSV/SARS-CoV-2/GFP wt_2E1_ populations were incubated with each of the nanobodies at a nanobody concentration that was 10x - 100x the IC50, to neutralize susceptible variants. Then the nanobody-virus mixture was plated on 293T/ACE2#22 cells, and neutralization resistant variants thereby selected and amplified by virus replication. Individual viral escape variants were then isolated by limiting dilution, amplified and their sensitivity to neutralization by the selecting nanobody compared to the sensitivity of the starting rVSV/SARS-CoV-2/GFP wt_2E1_ virus. We thus identified 35 rVSV-SARs-CoV-2/GFP mutants that exhibited resistance to each of the 20 nanobodies tested, selected on the basis of their high affinities, neutralization activities, and epitope coverage (**Suppl. Table 7**). For some of the less potent non-RBD epitope nanobodies, we used dimeric or trimeric forms of the nanobodies, but in each case the selected viral isolates exhibited resistance to monomeric, dimeric or trimeric forms. While some of the mutations that arose in the selection experiments were likely passenger mutations (**Suppl. Table 7**), a number of the mutations clustered on the Spike surface close to each other on RBD (**Fig. 7**) (Muecksch et al., 2021; Wang et al., 2021c; Weisblum et al., 2020). Some of the most potently neutralizing nanobodies selected resistant mutations at the same positions (e.g. E484K) as those selected by potent neutralizing antibodies that have been cloned from SARS-CoV-2 convalescents and vaccine recipients, confirming that the ACE2 binding site is a point of particular vulnerability for potent neutralization. Additionally, however, other nanobodies selected mutations that have not previously been encountered in human antibody selection experiments (**Suppl**. **Table 7**).

**Figure 7.**
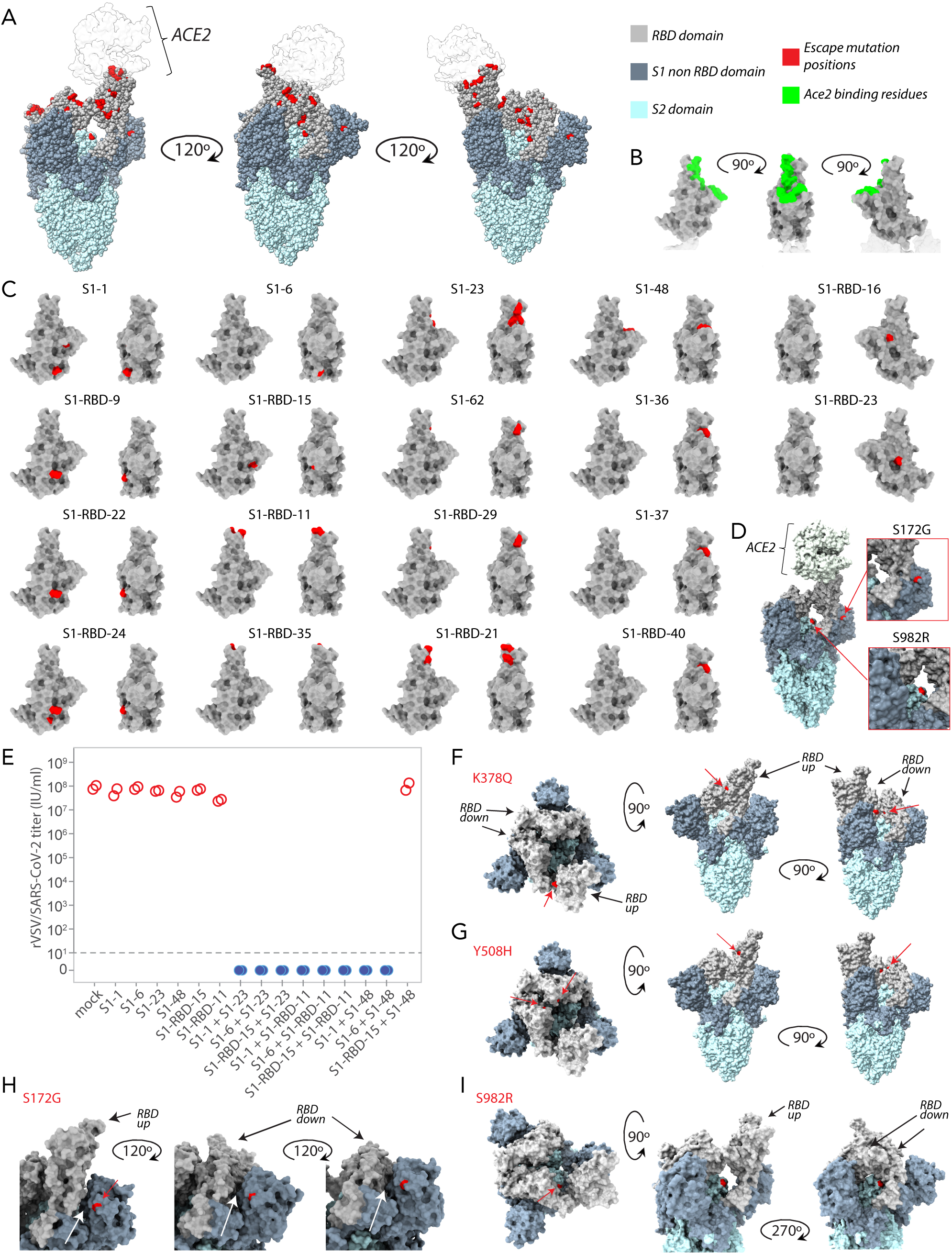
Mapping of Spike substitutions in rVSV/SARS-CoV-2/GFP escape mutants obtained in the presence of the corresponding nanobody. (A) Mapped on to the structure of SARS-CoV2 Spike trimer in complex with one ACE2 molecule (PDB ID 7KNB, used for all SARS-CoV2 Spike trimer representations) is the position of neutralization-resistant amino acid substitutions (in red), also known as ‘escape mutants’ that were generated in response to cultivation of rVSV/SARS-CoV-2/GFP in the presence of each nanobody, and were subsequently shown to confer resistance to the same nanobody. (B) Structure of the SARS-CoV-2 RBD (PDB ID 6M0J) showing the positions of amino acid residues (in green) that form the ACE2 binding site, for reference. (C) Structure of the SARS-CoV-2 RBD (PDB ID: 6M0J) showing the positions (in red) of the location of substitutions that confer resistance for each nanobody tested in two orientations 90° apart. For structure pairs of S1-RBD-16 and S1-RBD-23 escape mutants, the rotation is 90° from the structure to the left in the pair. (D) The location of two key non-RBD escape mutants S172G and S982K resulting from assays performed with an anti-S1 non RBD (S1-49) and anti-S2 (S2-10) nanobody respectively. (E) Infectious rVSV/SARS-CoV-2/GFP yield (IU/ml) following two passages in the presence of the indicated individual nanobodies or nanobody combinations, at 100x IC50 of the individual nanobodies, or 50 x IC50 of each of the nanobodies in the combinations. Each data point represents an independent titer measurement. Red open circles represent virus escapes while blue circles represent nanobody combinations for which no escapes (titer = 0) were detected. The location of two escape mutants K378Q (F) and Y508H (G) mapped onto SARS-CoV2 Spike trimer for the two corresponding nanobodies S1-RBD-9 and S1-RBD-15 respectively, revealing an exposed putative nanobody binding site on RBD when in the “up” position that is hidden when RBD is in the “down” position. (H) A close up of escape mutant S172G on each monomer of SARS-CoV2 Spike trimer revealing a larger crevice between the NTD of Spike S1 and RBD when the RBD is in the “down” position compared to the “up” position. (I) Three orientations of SARS-CoV2 Spike trimer revealing the position in all three orientations of escape mutant S982R revealing the putative binding site for nanobody S2-10 is accessible regardless of whether the RBD is the “up” or “down” position.

Beyond their enhanced combined activities (above), nanobody cocktails are expected to be resistant to escape (Baum et al., 2020; Gasparo et al., 2021; Weisblum et al., 2020). As proof of principle, we generated sets of two-nanobody cocktails by combining specific nanobodies that selected spatially distinct resistance mutations on the RBD (**Fig. 3A**). When rVSV/SARS-CoV2/GFP was passaged in the presence of the single nanobodies, resistant mutants were rapidly selected, as before. Indeed, the yield of infectious virus obtained after two passages in the presence of the single nanobody was nearly indistinguishable from that when rVSV/SARS-CoV2/GFP was passaged in the absence of nanobodies. In contrast, when nanobodies were combined in cocktails containing two nanobodies, at the same total concentration as was used for the individual nanobodies, in 8 out of 9 cases, no infectious rVSV/SARS-CoV2/GFP was recovered after two passages (**Fig. 7E**). In the ninth case in which S1-48 and RBD-15 were combined and virus was still recovered, sequence analysis revealed that this virus contained two amino acid substitutions, F490V and Y508H in the RBD. These substitutions were similar or identical to the individual substitutions found in the selection experiments with the single S1-48 and S1-RBD-15 nanobodies, which gave escape variants carrying the substitutions F490S and Y508H respectively (**Suppl. Table 7**). These results show that simply combining two nanobodies imposed the requirement for a minimum of two amino acid substitutions to confer resistance to the nanobody cocktail, greatly elevating the genetic barrier for escape. Such mixtures or derived multimers may represent powerful escape resistant therapeutics, and even more escape resistance should be possible by the use of three or more carefully chosen nanobodies in cocktails or multimers.

### Insights into the Mechanism of Nanobody Binding and Neutralization

Collectively epitope mapping, neutralization data and escape data can be used to deduce nanobody binding sites and speculate on mechanisms by which they inhibit the virus. For each nanobody, its escape mutants cluster around a highly restricted area on Spike that we interpret as corresponding to being part of its binding epitope. Overall, despite having raised nanobodies against S2, non-RBD S1 and RBD, neutralization activity and their corresponding escape mutants are distributed mainly over the ACE2-facing side of Spike (**Figs. 2A, 7A**); only ~20% of anti-S2 nanobodies and ~60% of non-RBD anti-S1 nanobodies are neutralizing, whereas ~80% of anti-RBD nanobodies are neutralizing with many escape mutants highly concentrated on the receptor-binding motif (RBM), the region of RBD that interacts directly with ACE2 and is most lightly glycosylated (Shajahan et al., 2020; Watanabe et al., 2020). This neutralization bias reflects the most obvious mechanism of viral inhibition, namely, blocking binding of Spike’s RBD domain to ACE2 on host membranes to preclude viral fusion, but the non-RBD based neutralization also underscores that other important mechanisms for viral inhibition exist.

The escape mutants for nanobodies that bind RBD are collectively spread across the entire surface of that domain, indicating that we are comprehensively exploring its available epitope space (**Fig. 7**). A subset of these nanobodies bind sites that interfere with ACE2 binding, preventing the virus from initial binding to its host cell (Starr et al., 2020; Walls et al., 2020b; Wrapp et al., 2020b). Even here, more than one category of inhibition exists. Nanobodies with epitopes that appear proximal to the ACE2 binding site, such as S1-1 and S1-RBD-15, may inhibit the virus by sterically hindering ACE2, such that their inhibitory mechanism may be via direct competition. Nanobodies, such as S1-23, S1-48, and S1-RBD-11, whose epitopes appear to directly overlap with the ACE2 binding site, may similarly competitively inhibit ACE2 binding, but may also mimic ACE2 binding and catalyze Spike trimer rearrangements that prematurely convert spike into a post-fusion state suppressing viral fusion. Combinations of these mechanisms can lead to strong synergistic effects as seen with the pairs S1-23 and S1-1, S1-23 and S1-RBD-15 (**Fig. 6B,C**). Based on the escape residue positions, both S1-1 and S1-RBD-15 bind to a similar site on one side of RBD, whereas S1-23 binds an adjacent and non-overlapping surface; other nanobodies with similar escape mutations likely also fall into one of these two binding classes (**Fig. 3C**, **Fig. 6**, **Fig. 7**). The ability of these nanobodies as pairs to sandwich RBD over the ACE2 binding site, may explain why such pairs are so strongly synergistic (**Fig. 6**). Binding of a nanobody to either region (**Fig. 6**) is expected to stabilize the otherwise “up”-”down” fluctuating RBD in its “up”, ACE2-engaging, position (**Fig. 7F,G**) (Bracken et al., 2021; Schoof et al., 2020a; Xiang et al., 2020). This can have three effects, all of which potentially promote nanobody synergy: first, it will increase the effective on-rate for Spike trimer to that measured for monomer - and therefore, make it easier for nanobodies from the other class to bind and inhibit; second, by stabilizing this “up” for any one of the three RBDs in each Spike trimer, it destabilizes the “down” position for the remaining two RBDs, again making nanobody binding from the second class more likely; and third, the “up” position exposes additional nanobody epitopes that would otherwise be buried (**Fig. 7**) (Sun et al., 2021; Xiang et al., 2020).

It is less clear how nanobodies with RBD epitopes very distal from the ACE2 binding site neutralize the virus. For example, neutralizing nanobodies S1-RBD-9 and S1-RBD-22, bind their overlapping epitopes >70 Å away from the ACE2 binding site and non-overlapping with the S1-1 and S1-23 epitopes (**Fig. 7C**). We hypothesize that these nanobodies can only bind when one RBD is “up”, but binding to the exposed site may block additional RBDs from moving to the up position; or, binding in this position at the base of the RBD and between S1 and S2 could destabilize the trimer, as has been proposed for other nanobodies (Sun et al., 2021). Interestingly, nanobodies sharing similar epitope bins as S1-RBD-9, such as S1-RBD-34, S1-RBD-19, S1-RBD-25, S1-RBD-32, and S1-RBD-36 do not neutralize - suggesting that regardless of the precise mechanism, the neutralization activity represented by S1-RBD-9 is specific to a very localized region on RBD.

A previously unstudied class of nanobodies are those that bind non-RBD domains of S1; we have 16 such nanobodies, 8 of which neutralized the virus. Mapping these via escapes proved challenging; however, the use of homodimers enhanced neutralization activity (above) allowing us to locate the epitope of S1-49 to the NTD of S1 (**Fig. 7D & H**). The position of the escape mutant at S172G suggests that a possible neutralization mechanism is one in which nanobody binding the larger crevice formed between the NTD and RBD when the latter is in the “down” position locks the RBD trimer in the “down” position to inhibit ACE2 binding (**Fig. 7H**). However, notably, human monoclonal antibodies specific to the NTD have been shown not to compete with ACE2 binding, and are instead proposed to inhibit viral infection by blocking membrane fusion, interaction with a different receptor, or proteolytic activation of Spike (McCallum et al., 2021). It remains to be determined if these mechanisms of neutralization hold for our nanobodies that bind non-RBD domains of S1, or if S1-49 suppresses ACE2 binding. The numerous human monoclonals that neutralize virus by binding outside of the RBD, and their yet to be discovered orthogonal mechanisms of neutralization, emphasize the potential, and need for further characterization, of our large repertoire of nanobodies.

The S2 domain is also a prime, but largely unexplored, therapeutic target (Elshabrawy et al., 2012; Shah et al., 2021). Here, we present the first neutralizing nanobodies that bind to S2 (**Fig. 2**, **Fig. 7I**). Some monoclonal antibodies that target S2 have been identified and shown to have neutralizing activity, but to our knowledge none have been structurally mapped (Andreano et al., 2021; Li et al., 2020; Poh et al., 2020; Song et al., 2020; Wang et al., 2021a). Because S2’s function is primarily membrane fusion rather than receptor binding, the nanobodies’ neutralization mechanisms must differ from those discussed above. For example, mapping of escape mutants of S2-10 (Fig. 7D) indicates binding at S982 (S982R) of spike, positioned at the end of the highly conserved heptad repeat 1, within a region of the S2 that undergoes large dynamic changes as the protein adopts a post fusion conformation; this suggests that S2-10 may restrict this conformational change, thereby inhibiting viral fusion (Cai et al., 2020; Pierri, 2020; Turonova et al., 2020; Walls et al., 2020b). Notably, the region proximal to S982 appears accessible through a ~30 Å portal, even in the prefusion form with the RBDs in the down position. This is a size not inconsistent with the binding face of a diminutive nanobody but likely inaccessible to conventional antibodies, as has been suggested by others (Xu et al., 2021).

The dimeric nature of conventional antibodies can introduce ambiguities regarding the mechanisms of neutralization, because they can operate either as individual or pairwise binders. In the latter case, they may operate, for example, by aggregation (Thomas et al., 1986), increased avidity, enhanced steric hindrance via the larger binding entity, or by simultaneously binding and locking two separate moieties within a viral particle. Nanobodies, as monomeric proteins, can provide a unique opportunity to differentiate between these possible mechanisms. In some cases, e.g., S1-7 and S1-25 (which are non-neutralizing as monomers), dimerization does not convert them into neutralizers. In other cases, dimerization and trimerization can engender several fold to orders of magnitude increase in neutralization potency (e.g., S1-RBD-35 and S1-23 respectively) (**Suppl. Table 3**). We even have a curious case where a nanobody such as S2-7 that is essentially non-neutralizing as a monomer becomes strongly neutralizing upon dimerization (**Fig. 4**). In this latter case, aggregation is a possible contributory mechanism, both between virions - which would lower effective virion concentration - or within a virion, with adjacent Spike trimers being crosslinked to each other, inhibiting their function. Although a tremendous range in neutralization improvements by oligomerization is observed both by ourselves and others, there is likely a limit to how much improvement can be induced by oligomerization as e.g. the trimers of S1-23 and S1-RBD-35 do not show a similar fold improvement as to what was observed for the monomer to dimer transition (Koenig et al., 2021; Ma et al., 2021; Schoof et al., 2020a; Xiang et al., 2020; Xu et al., 2021).

### Perspectives

The data presented here demonstrate the power of raising large and diverse repertoires of nanobodies against the entire ectodomain of SARS-CoV-2 Spike to maximize the likelihood of generating potent reagents for prophylactics and therapeutics. Our escape experiments support the idea that the current circulating variants are not yet necessarily exploring the full potential of the virus to escape our current and emerging therapeutic arsenals, and that even if antibodies or nanobodies are resistant to the current variants, they will not necessarily be resistant to variants as they continually emerge. However, we show that judicious choice of nanobody combinations that can synergize and have orthogonal and complementary neutralization mechanisms have the potential to result in broadly neutralizing reagents that are resistant to viral escape. Collectively, this large and readily modifiable repertoire promises to be of great value as therapeutics in the face of evolving variants, complementing vaccines, drugs and single epitope reagents, and guarding against single molecule failure in human trials.

## ACKNOWLEDGEMENTS

This work was supported by the G. Harold and Leila Y. Mathers Charitable Foundation, the Robertson Therapeutic Development Fund, the Jain Foundation, P41 grant NIH P41 GM109824 (J.D.A., B.T.C., M.P.R., A.S.), NIH R01AI501111 (P.D.B.), NIH R01AI078788 (T.H.), NIH U19AI125378 (J.S.B) and NIH K24AI150991 (J.S.B). We are very grateful to Sonya Paske and the staff at Capralogics, Inc. for raising antisera; the High-Throughput and Spectroscopy Resource Center (HTSRC), Rockefeller University, and especially Lavoisier Ramos-Espiritu for technical and data analysis support; Brenda Watt from Refeyn Ltd for technical and data analysis support; Jason Wendler and Maxwell Neal for help with statistical analyses; Tom Gallagher and Enya Qing, Loyola University Chicago for the kind gift of the SARS-CoV-1 and SARS-CoV-2 Spike expression plasmids; Andrew McGuire and Leonidas Stamatatos (Fred Hutchinson Cancer Research Center) for their kind gift of the SARS-CoV-2 20H/501Y.V2 Spike variant expression plasmid; and the rest of the Aitchison, Chait and Rout laboratories for technical and intellectual support.

---

**Supplementary Figure 1. Binding of nanobody candidates to immobilized antigen.**

All nanobody candidates identified by (a) S1, (b) S1-RBD, or (c) S2 purification were expressed in bacterial periplasm, which was bound to the respective immobilized antigen protein. After washes, loaded input (L) and elution (E) samples were analyzed by Coomassie stained SDS-PAGE. Positive binders (blue) displayed a nanobody band in the elution, while negative candidates (gray) had none.

**Supplementary Figure 2. Quantified antigen binding of nanobody candidates.**

All nanobody candidates were expressed in bacterial periplasm, which was bound to immobilized S1, S1-RBD, or S2 antigen protein. Bound nanobody was quantified by Coomassie staining after SDS-PAGE. Binding intensity against each antigen was normalized to the maximum observed binding among all nanobodies. Candidates with >20% maximum activity (blue) were selected for follow up, while others (grey) were generally discarded.

**Supplementary Table 1. S1 nanobody characterization.**

Nanobodies against S1 were determined to bind RBD or non-RBD epitopes by their affinity for recombinant full-length S1 and/or S1 RBD protein. Binding kinetics against these two recombinant proteins were determined by SPR, with on rates, off rates, and K_D_s determined by Langmuir fits to binding sensorgrams unless otherwise noted. Nanobody melting temperatures were determined by DSF. Nanobodies were assayed for neutralization activity against a SARS-CoV-2 Spike pseudotyped HIV-1 virus (PSV), with IC50s calculated from neutralization curves. Standard error of the mean (s.e.m.) is reported where replicates were available.

**Supplementary Table 2. S2 nanobody characterization.**

Binding kinetics of S2 nanobodies were determined by SPR using recombinant S2 protein, with on rates, off rates, and K_D_s determined by Langmuir fits to binding sensorgrams unless otherwise noted. Nanobody melting temperatures were determined by DSF. Nanobodies were assayed for neutralization activity against a SARS-CoV-2 or SARS-CoV-1 Spike pseudotyped HIV-1 virus (PSV), with IC50s calculated from neutralization curves. Standard error of the mean (s.e.m) is reported where replicates were available.

**Supplementary Table 3. Characterization of oligomerized Spike nanobodies.**

Nanobody monomers, dimers, or trimers were assayed for neutralization activity against a SARS-CoV-2 Spike pseudotyped HIV-1 virus (PSV), with IC50s calculated from neutralization curves. Standard error of the mean (s.e.m) is reported where replicates were available. Epitopes were determined by relative affinity for recombinant S1 or S1 RBD protein.

**Supplementary Table 4. Nanobody binding activity against Spike S1 variants.**

Binding kinetics against wild-type Spike S1 or two variants of concern were determined by SPR, with on rates, off rates, and K_D_s determined by Langmuir fits to binding sensorgrams.

**Supplementary Table 5. Nanobody neutralization activity against Spike variants.**

Nanobodies were assayed for neutralization activity against a pseudotyped HIV-1 virus (PSV) expressing SARS-CoV-2, SARS-CoV-1, or SARS-CoV-2 501Y.V2 Spike, with IC50s calculated from neutralization curves. Standard error of the mean (s.e.m) is reported where replicates were available.

**Supplementary Table 6. DSS-Crosslinked Nanobody-RBD Peptides Used for Modeling.**

S1-1, S1-23, and S1-RBD-15 nanobodies were bound to RBD and crosslinked with DSS (disuccinimidyl suberate). Crosslinked complexes were excised from SDS-PAGE gels, reduced, alkylated, and digested with either trypsin or chymotrypsin. Peptides were extracted and analyzed by mass spectrometry. Crosslinked peptides (listed) and residues (indicated by asterisk) were identified using pLink, and spectra were manually validated to eliminate false positives.

**Supplementary Table 7. Nanobody neutralization of rVSV/SARS-CoV-2 and selected resistant mutants.**

Neutralization assays carried out using rVSV/SARS-CoV-2 and 293T/ACE2cl.22 target cells with the denoted nanobodies their escape mutants (variants) and IC50s are listed.

## METHODS

### Antigens

Recombinant Fc-tagged SARS-CoV-2 Spike S1 and S2 proteins purified from HEK293 cells were used for llama immunization (The Native Antigen Company REC31806 and REC31807). For affinity isolation, binding screens, SPR analysis, and mass photometry, recombinant Spike S1-His, untagged RBD, or S2-His proteins expressed in HEK293 (S1 and RBD) or insect cells (S2) were used (Sino Biological 40591-V08H, 40592-VNAH, and 40590-V08B). Native mass spectrometry (Olinares et al., 2016; Olinares et al., 2021) was used to confirm the quality of these proteins, and were found to be glycosylated, with S1 and S2 observed to be heavily glycosylated (at least 10 kDa of attached glycans). RBD was observed to be monomeric, S1-His likely monomeric, and S2-His a mix of monomer and trimer.

### Immunization and Isolation of VHH Antibody Fractions

Two llamas, Marley (9 year old male) and Rocky (5 year old male) were immunized with recombinant SARS-CoV2 Spike S1 and SARS-CoV2 Spike S2 expressed in HEK293 cells as Fc fusion proteins. Llamas were injected subcutaneously with 0.25 mg of each antigen with CFA, then boosted with the same amounts with IFA at intervals of 14, 7, 21, and 10 days. Serum bleeds and bone marrow aspirates were obtained 9 days after the final injection. From the production serum bleeds, HCAb fractions of IgG were obtained by purification with immobilized Protein A and Protein G as previously described (Fridy et al., 2014). Residual light-chain containing IgG was removed from this fraction by incubating with 25 µl of 10 mg/ml Protein M-Sepharose per mg of HCAb (Grover et al., 2014). After a 30 min. incubation, the HCAb flow-through was collected. 12 mg of this HCAb fraction was then incubated with Sepharose-conjugated recombinant SARS-CoV2 Spike S1-His, RBD, or S2-His protein. This resin was washed with 1) 20 mM sodium phosphate, pH 7.4 + 500 mM NaCl; 2) 2 M MgCl_2_ in 20 mM Tris, pH 7.5; 3) PBS + 0.5% Triton X-100; and 4) PBS. The resin was then resuspended in a 200 μl solution of 2 U/µl IdeS enzyme (Genovis) in PBS, and digested for 3.5 hours at 37°C on an orbital shaker. The resin was then washed with 1) PBS 2) PBS plus 0.1% Tween-20 3) PBS. Bound protein was eluted by incubating 10 min. at 72°C in 1x NuPAGE LDS sample buffer (Thermo Fisher). The samples were reduced with DTT and alkylated with iodoacetamide, then run on a 4-12% Bis-Tris gel. Bands at ~15 kDa and ~20 kDa corresponding to digested V_H_H region were then cut out and prepared for MS.

### RT-PCR and DNA Sequencing

Bone marrow aspirates were obtained from immunized llamas concurrent with production serum bleeds. Bone marrow plasma cells were isolated on a Ficoll gradient using Ficoll-Paque (Cytiva). RNA was isolated from approximately 3-4 x 10^7^ cells using TRIzol reagent (Thermo Fisher), according to the manufacturer’s instructions. cDNA was synthesized using SuperScript IV reverse transcriptase (Thermo Fisher). A PCR was then performed with V_H_H IgG specific primers and Deep Vent polymerase (New England Biolabs). Forward primers 6N_CALL001 5′-NNNNNNGTCCTGGCTGCTCTTCTACAAGG-3′ and 6N_CALL001B 5′-NNNNNNGTCCTGGCTGCTCTTTTACAAGG-3′ target the leader sequence, (Conrath et al., 2001) while reverse primers 6N_VHH_SH_rev 5′-NNNNNNCTGGGGTCTTCGCTGTGGTGC-3′ and 6N_VHH_LH_rev 5′-NNNNNNGTGGTTGTGGTTTTGGTGTCTTGGG-3′ target short and long hinge sequences at the 3′ side of V_H_H. Primers included 6 random bases (N) to aid cluster identification. The approximately 350-450 bp product of this reaction was gel purified, then ligated to Illumina adaptors before library preparation using Illumina kits, before MiSeq sequencing using 2 x 300 bp paired end reads.

### Identification of Nanobodies by Mass Spectrometry

Trypsin (Roche) or chymotrypsin (Promega) solution were added to previously reduced, alkylated, dehydrated, diced and destained gel pieces at ~1:4-3:1 enzyme to substrate mass ratios and allowed to rehydrate for 10 min on ice. 45 ml of digestion buffer (trypsin: 50 mM ammonium bicarbonate, 10% acetonitrile, chymotrypsin: 100 mM Tris pH 7.8, 10 mM CaCl_2_) was then added, and samples were incubated for 6 h at 37 °C (trypsin) or 25 °C (chymotrypsin). Supernatant was then removed from gel pieces, and transferred to a new tube. 150 ml of a 1.67% FA, 67% ACN, 0.05% TFA solution were added to gel pieces, and shaken at 4°C for ~6h. Supernatant was removed from the gel pieces, transferred to the tube with the previous supernatant, and placed in a speedvac until dry. Samples were resuspended in 5% formic acid, 0.1% TFA and stage tipped to remove salt.

Samples were analyzed with a nano-LC 1200 (Thermo Fisher) using a EASYspray PepMap RSLC C18 3 micron, 100 Å, 75 um by 15 cm column coupled to an Orbitrap Fusion Lumos Tribrid mass spectrometer (Thermo Fisher). An Active Background Ion Reduction Device (ABIRD, ESI Source Solutions) was used to reduce background. The Lumos was operated in data-dependent mode, and top intensity ions were fragmented by high-energy collisional dissociation (normalized collision energy 28). Ions with charge states 2-5 were selected for fragmentation. Monoisotopic precursor selection mode (MIPS) was used to improve charge state identification. Target resolution was 120,000 for MS. The quadrupole isolation window was 1.4, and the MS/MS used a maximum injection time of 250☐ ms with 1 microscan and a minimum intensity threshold of 1E2.

The initial identification of nanobody sequences was performed as described (Fridy et al., 2014) using the program LlamaMagic with a few added features (including being able to deal with chymotryptic proteolysis and to rank VHH by corresponding read counts in high-throughput sequencing data), where 23 MS datasets (concatenated from all MS acquisition data according to antigens, animal individuals, gel band positions and proteases) were independently searched. The results were filtered with criteria including read counts, uniqueness score, and quality and coverage of MS/MS fragments to generate a collection of high confidence nanobody sequences. A CDR3 network graph was created by connecting nodes (unique high confidence CDR3 sequences) by edges where a CDR3 pair has a Damerau-Levenshtein distance of no more than three, by using NetworkX 2.5 (https://networkx.org) and pyxDamerauLevenshtein (https://github.com/gfairchild/pyxDamerauLevenshtein). The diversity of nanobodies for screening was maximized by selecting CDR3 sequences from isolated components of the network graph, together with varying CDR3 lengths and animal individual origin.

### Cloning and Purification of Nanobodies

Nanobody sequences were codon-optimized for expression in *E. coli* and synthesized as gene fragments (IDT), incorporating BamHI and XhoI restriction sites at 5′ and 3′ ends, respectively. Nanobody sequences were then subcloned into pET21-pelB using BamHI and XhoI restriction sites as previously described (Fridy et al., 2014). pelB-fused nanobodies were expressed and purified using Arctic Express (DE3) cells (Agilent) as previously described, using TALON metal affinity resin (Takara) (Fridy et al., 2014).

Nanobodies to be oligomerized were ordered from IDT as minigenes incorporating at the 5′ end a SalI site followed by codon optimized sequence for the linker GGGGSGGGGSGGGGSGGGGS upstream of the start codon of the nanobody cDNA, and at the 3′ end of the nanobody the coding sequence a XhoI site was added. The minigene was cut with SalI and XhoI, the linker-nanobody insert was gel purified and ligated with the XhoI linearized recipient nanobody expression vector (pET21-pelB+nanobody). Restriction digests and sequencing was performed to identify 2x (dimer) and 3x (trimer) oligomers.

### Nanobody Screening

To validate nanobody candidates, pelB-fused nanobodies were expressed in 50 ml cultures of Arctic Express (DE3) cells, and the periplasmic fractions were isolated by osmotic shock as previously described (Fridy et al., 2014). Spike S1-His, RBD, or S2-His proteins (Sino Biological 40591-V08H, 40592-VNAH, and 40590-V08B) were conjugated to cyanogen bromide-activated Sepharose 4 Fast Flow resin (Cytiva) according to the manufacturer’s instructions, using 100 µg protein per mg of resin. Periplasm was incubated with 15 µl of the corresponding antigen-conjugated Sepharose for 30 min while rotating at room temperature. The resin was then transferred to a spin column and washed twice with buffer TBT-100 (20 mM HEPES pH 7.4, 100 mM NaCl, 110 mM KOAc, 2 mM MgCl_2_, 0.1% Tween 20). Bound protein was eluted with 1.2 x NuPAGE LDS sample buffer (Thermo Fisher) for 10 min. at 72°C, then reduced with 50 mM DTT (10 min at 72°C). Input and elution samples were separated by SDS-PAGE and Coomassie-stained bands were quantified using ImageJ software.

### Surface Plasmon Resonance (SPR)

*K_D_*s were determined via surface plasmon resonance experiments. Measurements were either taken on a Proteon XPR36 Protein Interaction Array System (Bio-Rad) or a Biacore 8k (Cytiva). Recombinant Spike S1, RBD and Spike S2 was immobilized at X ug/ml, Y ug/ml and Z ug/ml respectively using EDC/NHS coupling chemistry according to the respective manufacturer’s guidelines either on a ProteOn GLC sensor chip or a Series S CM5 sensor chip. All purified nanobodies in a final buffer containing 20 mM HEPES, 150 mM NaCl, 0.02% Tween, were prepared in 5-8 concentrations. For experiments performed on the Proteon XPR36, protein was then injected at 50 μl/min for 120 s, followed by a dissociation time of 600 s. Residual bound proteins were removed by regenerating the chip surface using Glycine pH 3 + 1M MgCl_2_. Data were processed and analyzed using the ProteOn Manager software. For experiments performed on the Biacore 8k, protein was injected at 60 μl/min for 120 s, followed by a dissociation time of either 1200 s or 2400 s. Residual bound proteins were removed by regenerating the chip surface using Glycine pH 2.5 + 1M MgCl_2_. Data was processed and analyzed using the manufacturer’s software.

### Differential Scanning Fluorimetry (DSF)

Nanobody melting temperatures (*T*_M_) were measured by differential scanning fluorimetry (DSF) using a CFX96 Real-Time PCR Detection System (Bio-Rad, Hercules, CA). A 96-well thin-wall PCR plate (Bio-Rad) was set up with each well containing 10-40☐μM of protein in 20☐mM HEPES, 150☐mM NaCl buffer (pH 7.5), 5 × SYPRO Orange dye (SigmaAldrich). Fluorescence variation was measured from 25°C to 95°C at a ramp rate of 0.5°C/5 sec. Excitation was between 515–535 nm and emission was monitored between 560–580 nm. *T*m was the transition midpoint value, calculated using the manufacturer’s software (Niesen et al., 2007).

### Lyophilization

Nanobodies in 20 mM HEPES, 150 mM NaCl, pH 7.4 at concentrations between 0.33 mg/ml and 0.63 mg/ml were snap-frozen in liquid nitrogen and dried in a speed-vac to replicate lyophilization conditions. Nanobodies were then reconstituted in ddH2O and characterized using SPR and DSF.

### Epitope Mapping

Biolayer interferometry for epitope binning. Epitope mapping studies were carried out using the Octet system (ForteBio, USA, Version 7) that measures biolayer interferometry (BLI). All steps were performed at 30 °C with shaking at 1300 rpm in a black 96-well plate containing 300 μl kinetics buffer (PBS; 0.2%BSA; 0.02%Sodium Azide) in each well. AMC-coated biosensors were loaded with mFc tagged RBD (SinoBio) at 40 μg/ml to reach > 1 nm wavelength shift following binding and washing. The sensors were then reacted for ~300 s with reference nanobodies and then transferred to kinetics buffer-containing wells for another 180 s. A new baseline was set, sensors were then reacted for 180 s with analyte nanobodies (association phase) and then transferred to buffer-containing wells for another 180 s (dissociation phase). Binding and dissociation were measured as changes over time in light interference after subtraction of parallel measurements from unloaded biosensors. Sensorgrams of analyte association/dissociation responses were analyzed using the Octet data analysis software 7.1 (Fortebio, USA, 2015). Analyte binding to mFc RBD were also measured in parallel to get response levels in the absence of the reference nanobodies.

Octet response values were used to compute a Pearson’s Correlation Coefficient for pairwise combinations of nanobodies using Pandas (McKinney, 2010) in Python 3.7.6 (https://www.python.org/). These coefficients were then used to hierarchically cluster the nanobodies and were visualized as a heatmap (Pedregosa et al., 2011).

The undirected unweighted network graph of Octet response values was constructed by treating each nanobody as a node, adding an edge to each measured pair of different nanobodies, and setting the maximum response value of a nanobody pair as an attribute to the edge, by using NetworkX 2.5 (https://networkx.org). The least responses of pair-wise nanobodies within all fully measured nanobody subsets were computed by iterating through all network cliques of size 2 - 14 by using NetworkX’s “find_cliques” function, and taking the minimum value of edge attributes within each clique. Network coefficients (average shortest path length, average clustering coefficient and small-world coefficient sigma) were computed using NetworkX’s “average_shortest_path_length”, “average_clustering” and “sigma” functions. Network visualization was created by using D3.js (https://d3js.org).

### Mass Photometry

Select nanobodies were binned using mass photometry (MP). Experiments were performed on a Refeyn OneMP instrument (Refeyn Ltd). The instrument was calibrated with a mix of BSA (Sigma-Aldrich), thyroglobulin (Sigma-Aldrich) and beta-amylase (Sigma-Aldrich). Coverslips (Thorlabs) and gaskets (Grace Bio-Labs) were prepared by washing with 100% IPA followed by ddH_2_O, repeated 3 times, followed by drying with HEPA filtered air. 12 μl of buffer was added to each well to focus the instrument after which 8 μl of protein solution was added and pipetted up and down to briefly mix after which movies/frame acquisition was promptly started. The final concentration in each experiment of recombinant Spike S1 monomer (Sinobiological) and each nanobody was 30 nM and between 25-40 nM respectively. Movies were acquired for 60 s (6000 frames) using AcquireMP (version 2.3.0; Refeyn Ltd) using standard settings. All movies were processed, analyzed, and masses estimated by fitting a Gaussian distribution to the data using DiscoverMP (version 2.3.0; Refeyn Ltd).

### Cell Lines

Vero E6 cells (ATCC) were cultured at 30 °C in the presence of 5% CO2 in medium composed of high glucose Dulbecco’s modified Eagle’s medium (DMEM, Gibco) supplemented with 5% (v/v) heat inactivated fetal bovine serum (VWR). Cell line authentication was provided by the American Type Culture Collection. Cell cultures for these experiments were not tested for mycoplasma contamination.

293T/17 and 293T-hACE2 (Crawford et al., 2020) cells (Life Technologies Cat# R70007, RRID:CVCL_6911) were cultured in DMEM (Gibco) supplemented with 10% FBS, penicillin/streptomycin, 10 mM HEPES, and with 0.1 mM MEM non-essential amino acids (Thermo Fisher).

### Production of SARS-CoV-1, SARS-CoV-2, SARS-CoV-2 Variant Pseudotyped Lentiviral Reporter Particles

Pseudovirus stocks were prepared using a modified protocol published by (Crawford et al., 2020; Qing et al., 2020). Briefly, pseudovirus stocks were prepared by cotransfecting 4.75 µg pHAGE-CMV-Luc2-IRES-ZsGreen-W (BEI catalog number NR-52516) (Crawford et al., 2020), 3.75 µg psPAX and 1.5 µg spike containing plasmid using lipofectamine 3000 (Thermofisher). 4 x 10^6^ cells were plated 16-24 h prior to transfection. 60 h post transfection pseudovirus containing media was collected, cleared by centrifugation at 1000 *g* and filtered through a 0.45 µm syringe filter to clear debris. 1 ml aliquots were frozen at −80 °C. Pseudovirus was titered by 3-fold serial dilution on 293T-hACE2 cells (Crawford et al., 2020), treated with 2 µg/ml polybrene (Sigma). Infected cells were processed between 52-60 h by adding equal volume of Steady-Glo (Promega) and firefly luciferase signal was measured using the Biotek Model N4 with integration at 0.5 ms.

### SARS-CoV-2 Pseudovirus Neutralization Assay

All periplasmic purified nanobodies were treated with Triton X-114 to remove any residual endotoxins so as to not have endotoxins contribute to the effective neutralization. 293-hACE2 cells were plated at 2500-4000 cells per well on 384 solid white TC treated plates. 3-fold serially diluted nanobodies (10 dilutions in total) were incubated with 40,000-60,000 RLU equivalents of pseudotyped SARS-CoV-2-Luc for 1 h at 37 °C. Mock treatment, and a sham treatment with LaM2 nanobodies (Fridy et al., 2014) that do not bind to Spike were included as negative controls while untreated wells were used to monitor background levels. The nanobody-pseudovirus mixtures were then added in quadruplicate to 293T-hACE2 cells along with 2 µg/ml polybrene (Sigma). Cells were incubated at 37 °C with 5% CO2. Infected cells were processed between 52-60 h by adding equal volume of Steady-Glo (Promega) and firefly luciferase signal was measured using the Biotek Model N4 with integration at 0.5 ms. Data were processed using Prism7 (graphpad) Sigmoidal, 4PL, X is log(concentration) Least squares fit to calculate the logIC50 (half-maximal inhibitory concentration). All nanobodies were tested at least 2 times and with more than one pseudovirus preparation.

### Nanobody Synergy

Experiments were performed as per our Pseudovirus Neutralization Assay. A robotic liquid handler was used to prepare 2D matrices of serial dilutions of two nanobodies and then mix these with SARS-CoV-2 pseudovirus for 1 h. After incubation with the virus, the mixture was overlaid on a monolayer of 293-hACE2 cells and left to incubate for 56 h. Luminescence was quantified by addition of Steady-Glo reagent and luciferase activity was read out on a Biotek Model N4 with integration at 0.5 ms. Data was processed using synergy software (Wooten and Albert, 2020).

### Structural Analysis

Integrative structural modeling proceeded through the standard four-stage protocol (Kim et al., 2018; Rout and Sali, 2019; Russel et al., 2012; Saltzberg et al., 2021), which was scripted using the *Python Modeling Interface* package, a library for modeling macromolecular complexes based on the *Integrative Modeling Platform* software (Russel et al., 2012), version develop-31a0ad09b4 (https://integrativemodeling.org). The RBD, spanning amino acids R319-F541 was represented using the crystal structure of the co-complex of ACE2 bound RBD (PDB ID: 6M0J (Lan et al., 2020)). Comparative models of the nanobodies S1-1, S1-23 and S1-RBD-15 were built from the crystal structure of the human Vsig4 targeting nanobody Nb119 (PDB ID: 5IML (Wen et al., 2017)) as template using MODELLER (Sali and Blundell, 1993), and their CDR3 regions were refined using MODELLER’s loop modeling algorithm (Fiser et al., 2000). To maximize the efficiency of structural sampling while avoiding severe information loss, the system was represented at a resolution of one bead per residue, and the RBD and all three nanobodies were treated as rigid bodies. For each nanobody, alternate binding modes were scored using spatial restraints enforcing receptor-ligand shape complementarity, crosslink satisfaction, proximity of CDR3 and FR3 (framework) regions on the nanobodies to escape mutant residues on the RBD, and the simultaneous vs. competitive binding of nanobody pairs to the RBD. With the RBD fixed in space, 2,500,000 alternate docked nanobody models were produced through 50 runs of replica exchange Gibbs sampling based on the Metropolis Monte Carlo algorithm, where each Monte Carlo step consisted of a series of random rotations and translations of rigid nanobodies. The initial set of models was filtered to select an ensemble of 26,400 good scoring models through a model validation pipeline as detailed (Rout and Sali, 2019; Saltzberg et al., 2021; Viswanath et al., 2017). This ensemble satisfied all of the 27 crosslinks (for S1-1, S1-23 and S1-RBD-15) used for modeling, i.e. the crosslink distances were within the imposed 28 Å, and four out of six escape mutant residues were within the imposed 8 Å maximum distance to CDR3 and FR3 regions on the nanobodies; two escape mutant residues (Y369 for S1-1 and Y508 for S1-RBD-15) were within 10 Å of their respective nanobodies. S1-1 and S1-RBD-15 were found to share parts of their binding region on the same face of the RBD, while S1-23 localized on the opposite face (**Fig. 6B** and **Fig. 6C**). The violations for two of the escape mutant distance restraints can be rationalized due to the structural uncertainties introduced by the rigid representations of both the receptor and the nanobodies.

### SARS-CoV-2 Stocks and Titers

SARS-Related Coronavirus 2, Isolate USA-WA1/2020, NR-52281, was deposited by the Centers for Disease Control and Prevention and obtained through BEI Resources, NIAID, NIH. Viral stocks were propagated in Vero E6 cells. All experimental work involving live SARS-CoV-2 was performed at Seattle Children’s Research Institute (SCRI) in compliance with SCRI guidelines for BioSafety Level 3 (BSL-3) containment. An initial inoculum was diluted in Opti-MEM (Gibco) at 1:1000, overlaid on a monolayer of Vero E6 and incubated for 90 min. Following the incubation the supernatant was removed and replaced with 2% (v/v) FBS in Opti-MEM medium. The cultures were inspected for cytopathic effects, which were prominent after 48 h of infection. After 72 h, infectious supernatants were collected, cleared of cellular debris by centrifugation and stored at −80 °C until use. Viral titers were determined by plaque assay using a liquid overlay and fixation-staining method, as described (Case et al., 2020; Mendoza et al., 2020). Briefly, serially diluted virus stocks were used to infect confluent monolayers of Vero E6 cells (~1.2 x 10^6^ cells per well) cultured in six-well plates. After a 90 min incubation the virus was removed and the cell monolayer overlaid with a medium composed of 3% (w/v) carboxymethylcellulose and 4% (v/v) FBS in phenyl-free Opti-MEM. 96 h post infection, the viscous carboxymethylcellulose medium was removed and the cells were washed once with Dulbecco’s phosphate buffered saline (DPBS; Gibco) before being fixed with 4% (w/v) paraformaldehyde in DPBS. After a 30 min incubation the fixative was removed and the cells were rinsed with DBPS before being stained with 1% (w/v) crystal violet in 20% (v/v) ethanol. Contrast was enhanced by successive washes with DPBS and clear plaques representing individual viral infections were visualized as spots lacking the stain. Plaques were enumerated by first identifying the dilution factor of the well containing 10-100 plaques. After counting the plaques, the average number of plaque forming units (pfus) from three experiments was used to determine the viral titer by dividing the average by the dilution factor and volume of virus delivered per well.

### Focus Forming Reduction Assay with Authentic SARS-CoV-2

Nanobody neutralization of infectious SARS-CoV-2 was performed using a focus forming reduction assay. Briefly, eight three-fold serial dilutions of nanobodies were incubated with ~7.5 × 10^4^ focus forming units of SARS-CoV-2 for one hour at room temperature. The mixture was then added to a confluent monolayer of Vero E6 cells or 293-ACE2 (Crawford et al., 2020) plated at ~1.5 × 10^5^ cells per well and seeded in 48-well plates. 24 h post infection, the cells were washed once with DPBS, trypsinized with 0.05% trypsin (Gibco) and fixed for 30 min with 4% paraformaldehyde in DPBS. After fixation, the cells were permeabilized with 1% (w/v) Triton X-100 (Sigma Aldrich) for 30 min. After permeabilization, the cells were incubated with a blocking buffer (1% (w/v) bovine serum albumin (Calbiochem) and 0.5% (w/v) Triton X-100 in DBPS) for 60 min, and then stained with primary anti-Spike CR3022 (Absolute Antibody) monoclonal antibodies (1:1000), and secondary anti-human IgG antibodies (1:2000) conjugated to Alexa fluor 488 (Invitrogen). Cells staining positive for Spike were measured by flow cytometry on a Becton Dickinson BD LSR II Special Order System Flow Cytometer With HTS Sampler. The percentage of Spike-positive cells from triplicate wells for each dilution was used to determine the half maximal inhibitory concentrations (IC50) using a parametric 1D Hill fitting algorithm with synergy (Wooten and Albert, 2020). A mock treatment, sham treatment with LaM2 nanobodies (Fridy et al., 2014), and untreated cells were used as controls.

### SARS-CoV-2 Neutralization in Primary Airway Epithelial Cell (AEC) Cultures

Assays with primary airway epithelial cell cultures were performed as described (Barrow et al., 2021). Briefly, bronchial AECs were obtained under study #12490 approved by the Seattle Children’s Institutional Review Board, with investigations carried out following the rules of the Declaration of Helsinki of 1975. AECs were differentiated for 21 days at an air–liquid interface (ALI) on 12-well collagen-coated Corning® plates with permeable transwells in PneumaCult™ ALI media (Stemcell™, Vancouver, BC, Canada). Differentiated AECs were treated with nanobodies diluted in PBS, or PBS alone for 60 min, the liquid was removed, and the AECs were then infected with SARS-CoV-2 at a MOI of 0.5. At 24 h intervals the cells were treated with nanobodies or PBS for 60 min. After 96 h of infection, SARS-CoV-2 viral replication was measured in AEC cultures by quantitative PCR, with triplicate assays of harvested RNA from each SARS-CoV-2-infected AEC donor cell line (Genesig® Coronavirus Strain 2019-nCoV Advanced PCR Kit, Primerdesign®, Southampton, UK). The concentration of RNA harvested from AECs was used to normalize the qPCR data and was measured on a spectrophotometer (Nanodrop®).

### rVSV/SARS-CoV-2 Neutralization Assays

Nanobodies were five-fold serially diluted and then incubated with rVSV/SARS-CoV-2/GFP wt_2E1_ or plaque purified selected variants for 1 h at 37 °C. The nanobody/recombinant virus mixture was then added to 293T/ACE2.cl22 cells. After 16 h, cells were harvested, and GFP positive cells quantified by flow cytometry. The percentage of GFP+ cells was normalized to that derived from cells infected with rVSV/SARS-CoV-2 in the absence of nanobodies. The half-maximal inhibitory concentrations (IC50) for the nanobodies were determined using four-parameter nonlinear regression (least squares regression method without weighting) (GraphPad Prism).

### Sequence Analyses

To identify putative nanobody resistance mutations, RNA was isolated from aliquots of supernatant containing selected viral populations or individual plaque purified variants using NucleoSpin 96 Virus Core Kit (Macherey-Nagel). The purified RNA was subjected to reverse transcription using random hexamer primers and SuperScript VILO cDNA Synthesis Kit (Thermo Fisher Scientific). The cDNA was amplified using KOD Xtreme Hot Start DNA 396 Polymerase (Millipore Sigma) flanking the Spike encoding sequences. The PCR products were gel-purified and sequenced using Sanger-sequencing.

### Selection of Virus Variants in the Presence of Nanobodies

For selection of Spike variants that were resistant to nanobodies, rVSV/SARS-CoV-2/GFP wt_2E1_ was passaged to generate diversity and populations containing 10^6^ infectious particles were used. The rVSV/SARS-CoV-2/GFP wt_2E1_ populations were incubated with dilutions of nanobodies (10 to 100× the IC50 excess) for 1 h at 37 °C. Then, the virus-nanobody mixtures were incubated with 5 × 10^5^ 293T/ACE2.22 cells in 6-well plates. Two days later, the cells were imaged and supernatant were harvested from cultures that showed evidence of viral replication (GFP-positive foci) or large numbers of GFP-positive cells. A 100 μl of the cleared supernatant was incubated with the same dilution of nanobodies and then used to infect 5 × 10^5^ 293T/ACE2.22 cells in 6-well plates, as before. rVSV/SARS-CoV-2/GFP2E1 were passaged in the presence combination of nanobodies two times before complete escape was evaluated. To isolate individual mutant viruses, selected rVSV/SARS-CoV-2/GFP wt_2E1_ populations were serially diluted in medium without nanobodies and individual viral variants isolated by visualizing single GFP-positive plaques at limiting dilutions in 96-well plates containing 1 × 10^4^ 293T/ACE2.22 cells. These plaque-purified viruses were expanded, and further characterized using sequencing and nanobody neutralization assays.

### Crosslinking Mass Spectrometry

Nanobodies and antigens were incubated together at a 2× molar excess of nanobody at RT for 10 min in 20 mM HEPES pH 7.4 and 150 mM NaCl. Crosslinker was then added to a final concentration of 5 mM bissulfosuccinimidyl suberate (BS3) or 1 mM disuccinimidyl suberate (DSS), and samples were crosslinked for 30 min at RT. Reactions were quenched, reduced and alkylated, and run on an SDS-PAGE gel. The band corresponding to the crosslinked nanobody-antigen complex was then excised from the gel and subjected to in-gel digestion with trypsin (Roche, 1μg, 4 h) or chymotrypsin (Roche, 0.5μg, 1.5 h).

Peptides were extracted and analyzed with a nano-LC 1200 (Thermo Fisher) with a EASYspray PepMap RSLC C18 3 micron, 100 Å, 75 μm by 15 cm column coupled to an Orbitrap Fusion Lumos Tribrid mass spectrometer (Thermo Fisher). An Active Background Ion Reduction Device (ABIRD, ESI Source Solutions) was used to reduce background. The Lumos was operated in data-dependent mode, and ions were fragmented by high-energy collisional dissociation (normalized collision energy 28). Separate LC runs were used to analyze the +3 and the +4 through +7 charge states. Higher charge species were prioritized for selection for fragmentation when analyzing the 4-7 species. Monoisotopic precursor selection mode (MIPS) was used to improve charge state identification. Target resolution was 30,000 for MS and 15,000 for MS/MS analyses. The quadrupole isolation window was 1.4, and the MS/MS used a maximum injection time of 800☐ms with 4 microscans and a minimum intensity threshold of 1E2. Data were then searched by pLink 2.3 (Chen et al., 2019) in order to identify cross-linked peptides. The mass accuracy in pLink was set to 10 p.p.m. for MS and 20☐p.p.m. for MS/MS. Cysteine carbamidomethylation was included as a fixed modification and methionine oxidation as a variable modification. For trypsin, up to three missed-cleavages were permitted. For chymotrypsin, the enzyme setting was “non-specific.” Spectra were manually checked to ensure correct identifications of crosslinked peptides.

