## Supplementary figures and images for "Nanobody Repertoires for Exposing Vulnerabilities of SARS-CoV-2"

### Supplementary Figure 1

#### Supplementary Figure 1

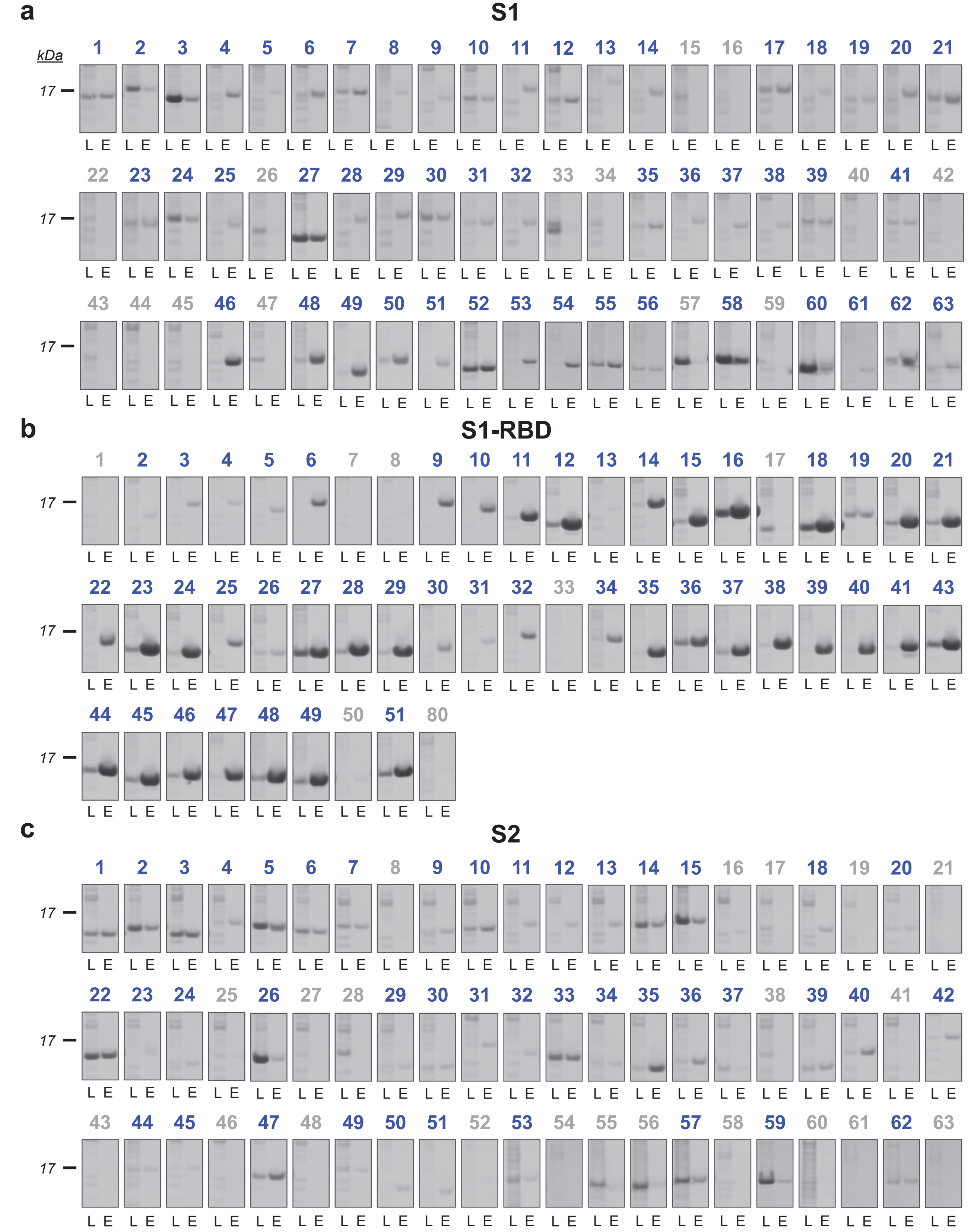

### Supplementary Figure 2

##### Supplementary Figure 2

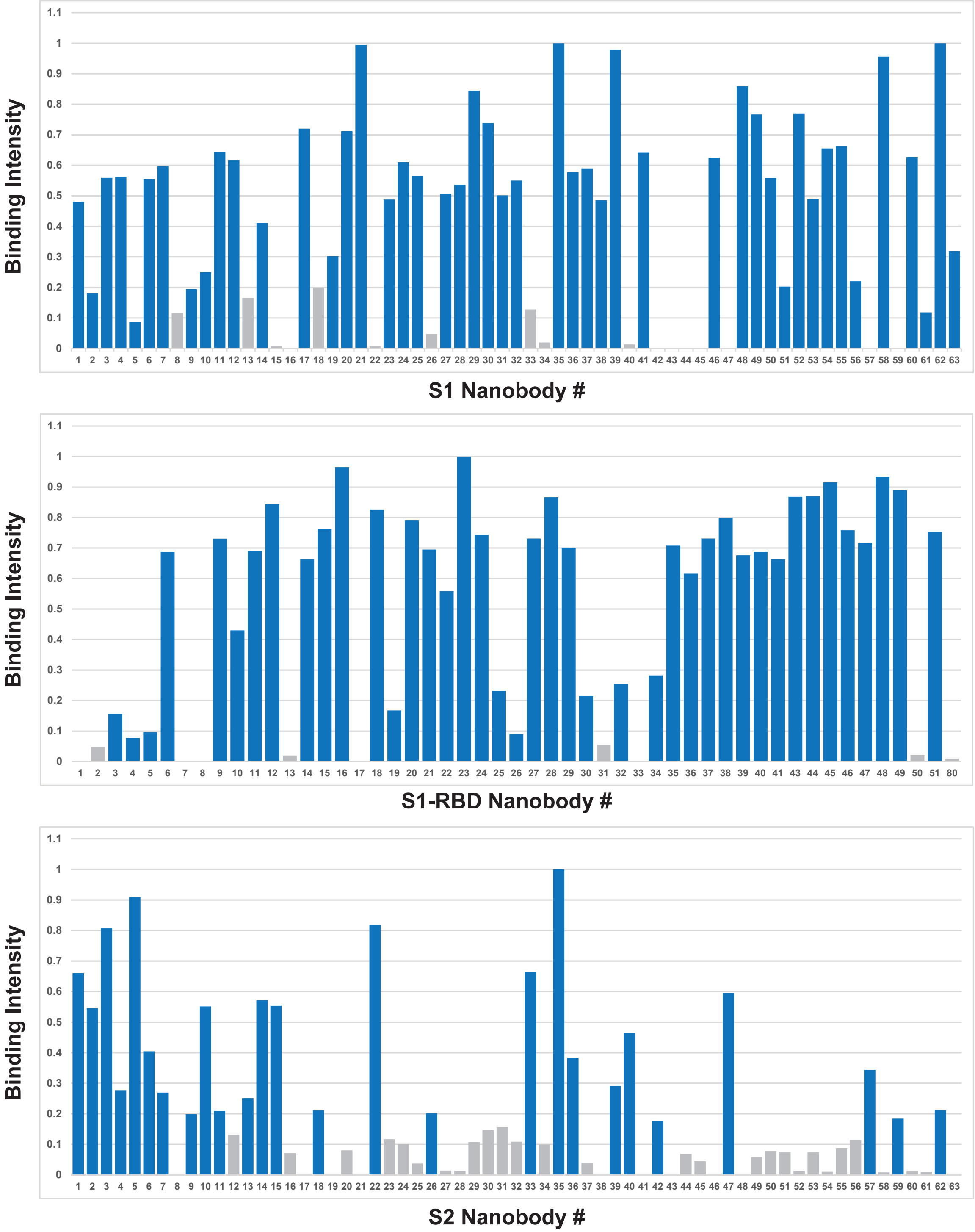
