## Supplementary Table 1 for "Nanobody Repertoires for Exposing Vulnerabilities of SARS-CoV-2"

| ID | Epitope | S1 | | | RBD | | | $T_m$ (ºC) | SARS-CoV-2 PSV IC50 (s.e.m) (nM) |
|---|---|---|---|---|---|---|---|---|---|
| | | $K_{on}$ ($M^{-1}s^{-1}$) | $K_{off}$ ($s^{-1}$) | $K_D$ (M) | $K_{on}$ ($M^{-1}s^{-1}$) | $K_{off}$ ($s^{-1}$) | $K_D$ (M) | | |
| S1-1 | RBD | 4.14E+05 | 2.98E-05 | 7.20E-11 | 6.50E+05 | 5.98E-07 | 9.20E-12 | 66.5 | 6.74 (1.03) |
| S1-2 | non-RBD | 1.59E+06 | 1.88E-03 | 1.18E-09 | No interaction detected | | | 66.5 | NA |
| S1-3 | non-RBD | 5.08E+05 | 4.32E-04 | 8.51E-10 | No interaction detected | | | 64 | 1030 (666) |
| S1-4 | RBD | 1.25E+06 | 1.06E-04 | 8.46E-11 | 1.26E+06 | 1.26E-04 | 1.37E-10 | 66 | 41.5 (3.68) |
| S1-5 | RBD | 1.33E+05 | 1.15E-03 | 8.61E-09 | – | | | 65.25 | NA |
| S1-6 | RBD | 1.02E+06 | 5.75E-04 | 5.65E-10 | 5.92E+05 | 3.69E-04 | 6.22E-10 | 65 | 56.1 (20.7) |
| S1-7 | non-RBD | 7.59E+05 | 9.90E-04 | 1.30E-09 | No interaction detected | | | 60.5 | NA |
| S1-9 | non-RBD | 9.51E+05<br>1.25E+05 | 4.28E-07<br>1.57E-04 | 4.50E-13[a]<br>1.25E-09 | – | | | 47.5 | NA |
| S1-10 | non-RBD | 8.35E+04 | 1.82E-03 | 2.19E-08 | – | | | 64 | NA |
| S1-11 | non-RBD | – | | | – | | | 60 | NA |
| S1-12 | RBD | 2.90E+05 | 8.92E-04 | 3.07E-09 | 2.33E+05 | 2.24E-04 | 9.63E-10 | 68 | NA |
| S1-14 | RBD | 1.08E+06 | 1.10E-03 | 1.02E-09 | 5.37E+05 | 7.99E-04 | 1.49E-09 | 57.5 | 135.8 (36.4) |
| S1-17 | non-RBD | – | | | – | | | 65 | 1271 (888) |
| S1-19 | RBD | 1.30E+06<br>3.55E+04 | 8.86E-03<br>2.41E-04 | 6.81E-09[a]<br>6.81E-09 | – | | | 64.5 | 139 (9.58) |
| S1-20 | RBD | 1.48E+07 | 4.37E-03 | 2.95E-10 | – | | | 69 | 51.8 (3.72) |
| S1-21 | RBD | 4.77E+06 | 1.58E-04 | 3.31E-11 | 1.22E+06 | 2.45E-04 | 2.00E-10 | 70.5 | 68.1 |
| S1-23 | RBD | 2.82E+06 | 4.91E-05 | 1.74E-11 | 1.09E+06 | 1.07E-04 | 9.78E-11 | 64 | 5.7 (2.163) |
| S1-24 | non-RBD | 6.49E+05 | 2.89E-04 | 4.45E-10 | No interaction detected | | | 71.5 | 868 |
| S1-25 | non-RBD | 2.15E+05 | 3.39E-05 | 1.57E-10 | No interaction detected | | | 58 | NA |
| S1-27 | RBD | 3.15E+06 | 4.52E-04 | 1.43E-10 | 2.89E+06 | 6.30E-04 | 2.18E-10 | 54 | 19.5 (4.90) |
| S1-28 | RBD | 1.38E+06 | 7.97E-04 | 5.76E-10 | 1.79E+06 | 1.03E-03 | 5.77E-10 | 66 | 66.0 (10.9) |
| S1-29 | RBD | 2.39E+05 | 1.01E-03 | 4.21E-09 | 1.73E+05 | 8.89E+04 | 5.12E-09 | 61.5 | NA |
| S1-30 | non-RBD | 6.21E+05 | 1.48E-03 | 2.38E-09 | No interaction detected | | | 57 | 330 |
| S1-31 | RBD | 2.17E+06 | 5.63E-04 | 2.59E-10 | 1.94E+06 | 9.37E-04 | 4.84E-10 | 72 | 78.7 (3.48) |
| S1-32 | non-RBD | 2.73E+05 | 4.66E-04 | 1.71E-09 | No interaction detected | | | 79 | NA |
| S1-35 | RBD | 2.46E+06 | 2.11E-05 | 8.60E-12 | 2.70E+06 | 9.77E-05 | 3.62E-11 | 70.5 | 12.5 (0.101) |
| S1-36 | RBD | 2.28E+06 | 3.92E-04 | 1.72E-10 | 7.87E+06 | 1.72E-03 | 2.18E-10 | 63 | 48.5 (21.1) |
| S1-37 | RBD | 4.03E+06 | 2.75E-04 | 6.82E-11 | 4.14E+06 | 2.09E-04 | 5.05E-11 | 65 | 7.54 |
| S1-38 | RBD | 5.34E+06 | 1.12E-03 | 2.10E-10 | – | | | 64 | 63.2 |
| S1-39 | RBD | 2.14E+06 | 8.11E-04 | 3.79E-10 | 1.68E+06 | 1.06E-03 | 6.30E-10 | 55 | 111 (4.03) |
| S1-41 | non-RBD | 8.73E+05 | 1.38E-03 | 1.58E-09 | No interaction detected | | | 62.5 | 732 |
| S1-46 | RBD | 1.68E+05 | 2.94E-04 | 1.75E-09 | 2.22E+05 | 1.70E-04 | 7.66E-10 | 68 | 312 (14.0) |
| S1-48 | RBD | 2.61E+06 | 6.22E-05 | 2.39E-11 | 1.66E+06 | 1.64E-04 | 9.85E-11 | 60.5 | 5.82 (0.832) |
| S1-49 | non-RBD | 1.94E+06 | 3.63E-03 | 1.87E-09 | – | | | 49, 74[c] | 356 (56.9) |
| S1-50 | non-RBD | 3.33E+05<br>3.34E-03 | 1.39E-02<br>3.94E-04 | 4.40E-09[b] | No interaction detected | | | 66 | 24,303 |
| S1-51 | RBD | 9.28E+04 | 4.22E-04 | 4.54E-09 | 3.77E+06 | 2.01E-03 | 5.33E-10 | 56 | 608 |
| S1-52 | RBD | 4.22E+05 | 3.13E-04 | 7.74E-09 | 4.53E+04 | 1.94E-04 | 4.36E-09 | 57.5 | 3,634 |
| S1-53 | RBD | 1.40E+06<br>2.36E+04 | 8.46E-03<br>2.19E-04 | 6.05E-09<br>9.27E-09 | – | | | 51.5 | 3,406 |
| S1-54 | RBD | 1.13E+06 | 6.58E-05 | 5.84E-11 | 2.55E+04 | 2.88E-04 | 1.13E-08 | 69 | 148 |

| | | | | | | | | | |
|---|---|---|---|---|---|---|---|---|---|
| S1-55 | RBD | 3.98E+06<br>3.53E+04 | 5.41E-03<br>5.31E-06 | 1.36E-09<br>1.51E-10 | 5.03E+05<br>1.84E+04 | 1.11E-02<br>1.82E-04 | 2.21E-08<br>9.89E-09 | 54.5 | 2,353 |
| S1-56 | RBD | 1.46E+04<br>4.45E-03 | 2.99E-03<br>7.90E-05 | 3.57E-09[b] | 2.21E+03 | 1.05E-04 | 4.73E-08 | 54 | NA |
| S1-58 | non-RBD | 5.73E+05 | 1.66E-04 | 2.90E-10 | | – | | 53.5 | 146 |
| S1-60 | non-RBD | 3.30E+05<br>4.61E+04 | 5.24E-06<br>3.67E-03 | 1.59E-11[a]<br>9.58E-08 | | – | | 62 | NA |
| S1-61 | RBD | 9.87E+05<br>8.23E+02 | 1.81E-02<br>1.10E-04 | 1.84E-08[a]<br>1.34E-07 | 4.46E+04 | 1.88E-04 | 4.21E-09 | 60 | NA |
| S1-62 | RBD | 2.68E+06 | 9.51E-05 | 3.54E-11 | 3.30E+06 | 6.30E-05 | 2.08E-11 | 71.5 | 4.95 |
| S1-63 | RBD | 1.09E+06<br>3.39E+04 | 1.12E-02<br>1.67E-04 | 1.02E-08[a]<br>4.94E-09 | 5.10E+04 | 2.23E-04 | 4.37E-09 | 65 | NA |
| S1-RBD-3 | RBD | 8.81E-05<br>7.36E+04 | 1.76E-02<br>1.13E-03 | 2.00E-08[a]<br>1.53E-08 | | – | | 72 | 384 (18.7) |
| S1-RBD-4 | RBD | 2.02E+06 | 1.64E-04 | 8.09E-11 | 2.83E+06 | 8.16E-04 | 2.89E-10 | 64.5 | 17.5 (1.98) |
| S1-RBD-5 | RBD | 1.94E+06 | 1.63E-04 | 8.38E-11 | 7.21E+06 | 1.05E-03 | 1.45E-10 | 64 | 34.5 |
| S1-RBD-6 | RBD | 1.55E+06 | 1.63E-04 | 1.05E-10 | 3.48E+06 | 1.13E-03 | 3.24E-10 | 66.5 | 77.2 (21.8) |
| S1-RBD-9 | RBD | | – | | 2.85E+05 | 1.23E-04 | 4.30E-10 | 69 | 522 |
| S1-RBD-10 | RBD | | – | | | – | | – | 52.9 |
| S1-RBD-11 | RBD | 2.22E+07 | 2.94E-04 | 1.32E-11 | 2.06E+07 | 4.06E-04 | 1.97E-11 | 65 | 13.5 (5.50) |
| S1-RBD-12 | RBD | | – | | 1.10E+04 | 3.39E-05 | 3.10E-09 | 67 | NA |
| S1-RBD-14 | RBD | | – | | 1.33E+04 | 3.34E-04 | 2.51E-08 | 65 | NA |
| S1-RBD-15 | RBD | 5.37E+06 | 1.50E-04 | 2.79E-11 | 7.52E+06 | 4.95E-04 | 6.58E-11 | 59.5, 80[c] | 5.29 (1.44) |
| S1-RBD-16 | RBD | | – | | 1.68E+04 | 6.25E-05 | 3.73E-09 | 61 | 79.2 (4.23) |
| S1-RBD-18 | RBD | 2.28E+06 | 6.25E-04 | 2.74E-10 | 4.43E+06 | 1.27E-03 | 2.87E-10 | 69.5 | 67.2 (1.92) |
| S1-RBD-19 | RBD | | – | | | – | | 60 | 3,902 |
| S1-RBD-20 | RBD | 2.37E+06 | 2.23E-04 | 9.43E-11 | 3.05E+06 | 7.91E-04 | 2.59E-10 | 49, 70[c] | 12.4 (1.06) |
| S1-RBD-21 | RBD | 3.50E+06 | 1.31E-03 | 3.73E-10 | 3.15E+06 | 1.71E-03 | 5.45E-10 | 48.5, 70.5[c] | 14.1 |
| S1-RBD-22 | RBD | 9.34E+05 | 2.28E-04 | 2.44E-10 | 9.24E+05 | 4.42E-04 | 4.78E-10 | 57.5 | 100 (0.078) |
| S1-RBD-23 | RBD | | – | | 2.89E+06 | 4.61E-05 | 1.59E-11 | 61 | 7.31 (0.418) |
| S1-RBD-24 | RBD | 1.61E+06 | 1.40E-03 | 8.65E-10 | 2.12E+06 | 1.22E-03 | 5.75E-10 | 46, 67[c] | 218 |
| S1-RBD-25 | RBD | | – | | 8.41E+04 | 1.16E-02 | 1.38E-07 | – | NA |
| S1-RBD-26 | RBD | 1.06E+05 | 4.58E-06 | 4.32E-11 | 2.15E+05 | 1.33E-05 | 6.19E-11 | 66 | 241 (81.4) |
| S1-RBD-27 | RBD | | – | | 6.19E+06 | 1.24E-02 | 2.00E-09 | 71 | 65.8 |
| S1-RBD-28 | RBD | 1.80E+06 | 4.27E-04 | 2.38E-10 | 1.80E+06 | 4.27E-04 | 2.38E-10 | 64.5 | 32.7 (3.07) |
| S1-RBD-29 | RBD | | – | | 5.36E+05 | 1.35E-03 | 2.51E-09 | 74 | 9.53 (1.04) |
| S1-RBD-30 | RBD | 2.15E+06 | 6.66E-05 | 3.10E-11 | 3.77E+06 | 4.82E-04 | 1.28E-10 | 65 | 25.0 (3.57) |
| S1-RBD-32 | RBD | | – | | 1.05E+05 | 7.90E-03 | 7.52E-08 | 65 | NA |
| S1-RBD-34 | RBD | | – | | 5.71E+04 | 4.88E-03 | 8.54E-08 | 64 | NA |
| S1-RBD-35 | RBD | 8.01E+05 | 1.68E-04 | 2.10E-10 | 1.33E+06 | 2.50E-04 | 1.88E-10 | 57, 68[c] | 12.3 (2.40) |
| S1-RBD-36 | RBD | | – | | | – | | 71 | NA |
| S1-RBD-37 | RBD | | – | | 3.60E+05 | 8.88E-04 | 2.47E-09 | 71 | 391 |
| S1-RBD-38 | RBD | | – | | 1.12E+06 | 9.84E-04 | 8.79E-10 | 68.5 | 84.6 (22.7) |
| S1-RBD-39 | RBD | | – | | 4.92E+05 | 7.77E-05 | 1.58E-10 | 67.5 | 77.8 |
| S1-RBD-40 | RBD | | – | | 7.47E+05 | 2.77E-05 | 3.71E-11 | 70 | 25.6 (5.88) |
| S1-RBD-41 | RBD | | – | | 4.37E+05 | 1.39E-04 | 3.17E-10 | – | 17.0 |
| S1-RBD-43 | RBD | | – | | 6.21E+05 | 1.82E-04 | 2.92E-10 | 68 | 33.6 (1.33) |
| S1-RBD-44 | RBD | | – | | 1.91E+05 | 6.97E-05 | 3.65E-10 | 57.5 | 93.4 |
| S1-RBD-45 | RBD | | – | | 4.43E+05 | 4.14E-05 | 9.30E-11 | 53 | 22.6 |

| | | | | | | | | | |
|---|---|---|---|---|---|---|---|---|---|
| S1-RBD-46 | RBD | – | | | 4.69E+05 | 5.79E-04 | 1.23E-09 | 75.5 | 48.0 |
| S1-RBD-47 | RBD | – | | | 2.11E+05 | 6.45E-04 | 3.06E-09 | 53.5 | 127 (11.6) |
| S1-RBD-48 | RBD | – | | | 1.05E+05 | 2.42E-04 | 2.30E-09 | 58, 63[c] | 54.5 |
| S1-RBD-49 | RBD | 3.24E+05 | 3.24E-04 | 1.00E-09 | 3.15E+05 | 5.34E-04 | 1.69E-09 | 66.5 | 37.6 |
| S1-RBD-51 | RBD | – | | | 3.77E+06 | 2.01E-03 | 5.33E-10 | 52, 61[c] | 70.9 (29.3) |

[a]Curves were fit to a heterogeneous ligand model. Respective $K_{on}$, $K_{off}$, and $K_D$ values are shown for each component.

[b]Curves were fit to a two-state reaction model. Respective $K_{on}$, $K_{off}$, and $K_D$ values are shown for each binding state.

[c]Two peaks were observed in the melting curve. $T_m$s for both are reported.

"–" - Not determined

NA - No activity
