## Supplementary Table 2 for "Nanobody Repertoires for Exposing Vulnerabilities of SARS-CoV-2"

| ID | $K_{on}$ ($M^{-1}s^{-1}$) | $K_{off}$ ($s^{-1}$) | $K_D$ (M) | $T_m$ (ºC) | SARS-CoV-2 PSV IC50 (s.e.m) (nM) |
|---|---|---|---|---|---|
| S2-1 | 6.32E+04 | 3.79E-04 | 6.00E-09 | 65.5 | NA |
| S2-2 | 1.26E+06 | 9.35E-05 | 7.41E-11 | 64.5 | 4,460 (902) |
| S2-3 | 2.62E+05 | 7.21E-05 | 2.76E-10 | 65 | 2,234 (751) |
| S2-4 | 2.44E+06 | 2.62E-04 | 1.08E-10 | 56 | NA |
| S2-5 | 9.35E+05 | 2.74E-04 | 2.93E-10 | 66 | NA |
| S2-6 | | – | | 61.5 | NA |
| S2-7 | 1.66E+06 | 9.36E-05 | 5.62E-11 | 61 | NA |
| S2-9 | 9.29E+05 | 2.32E-04 | 2.50E-10 | 64.5 | NA |
| S2-10 | 9.31E+04 | 3.13E-04 | 3.37E-09 | 59, 64.5[a] | 1,718 |
| S2-11 | 7.94E+06 | 1.12E-03 | 1.41E-10 | 69.5 | NA |
| S2-13 | 7.02E+05 | 1.05E-04 | 1.49E-10 | 64.5 | NA |
| S2-14 | 3.16E+06 | 1.28E-03 | 4.07E-10 | 72.5 | NA |
| S2-15 | | – | | 70 | NA |
| S2-18 | 1.63E+06 | 4.87E-04 | 2.99E-10 | 47, 54.5[a] | NA |
| S2-22 | | – | | – | NA |
| S2-26 | 4.45E+05 | 8.15E-05 | 1.83E-10 | 76.5 | NA |
| S2-33 | 3.68E+05 | 5.58E-05 | 2.33E-10 | 70 | NA |
| S2-35 | 2.36E+05 | 4.72E-05 | 2.00E-10 | 77 | NA |
| S2-36 | 4.39E+06 | 3.69E-04 | 8.41E-11 | 74 | NA |
| S2-39 | | – | | 58 | NA |
| S2-40 | 5.08E+04 | 7.16E-05 | 1.41E-09 | 69.5 | 1,712 (828) |
| S2-42 | 5.12E+05 | 3.77E-06 | 7.36E-12 | 69 | NA |
| S2-47 | 3.86E+05 | 1.14E-04 | 2.96E-10 | 40, 65[a] | NA |
| S2-57 | 2.33E+06 | 7.18E-04 | 3.08E-10 | 67 | NA |
| S2-59 | | – | | 37.5, 60[a] | NA |
| S2-62 | 1.65E+06 | 1.12E-04 | 6.81E-11 | 64, 77.5[a] | 12,979 |

[a]Two peaks were observed in the melting curve. $T_m$s for both are reported.

"–" - Not determined

NA - No activity
