## Supplementary Table 3 for "Nanobody Repertoires for Exposing Vulnerabilities of SARS-CoV-2"

| ID | Epitope | SARS-CoV-2 PSV IC50 (s.e.m) (nM) |
|---|---|---|
| S1-1 | RBD | 6.74 (1.03) |
| S1-1$_{dimer}$ | RBD | 5.09 |
| S1-1$_{trimer}$ | RBD | 5.87 |
| S1-23 | RBD | 5.69 (2.16) |
| S1-23$_{dimer}$ | RBD | 0.22 (0.054) |
| S1-23$_{trimer}$ | RBD | 0.089 (0.019) |
| S1-RBD-35 | RBD | 12.3 (2.40) |
| S1-RBD-35$_{dimer}$ | RBD | 0.15 (0.11) |
| S1-RBD-35$_{trimer}$ | RBD | 0.068 (0.043) |
| S1-3 | S1 non-RBD | 1030 (666) |
| S1-3$_{dimer}$ | S1 non-RBD | 429 |
| S1-30 | S1 non-RBD | 330 |
| S1-30$_{dimer}$ | S1 non-RBD | 18.3 |
| S1-7 | S1 non-RBD | NA |
| S1-7$_{dimer}$ | S1 non-RBD | NA |
| S1-7$_{trimer}$ | S1 non-RBD | NA |
| S1-17 | S1 non-RBD | 1271 (888) |
| S1-17$_{dimer}$ | S1 non-RBD | 2,144 |
| S1-49 | S1 non-RBD | 356 (56.9) |
| S1-49$_{dimer}$ | S1 non-RBD | 9.06 (1.20) |
| S1-49$_{trimer}$ | S1 non-RBD | 0.87 (0.075) |
| S2-7 | S2 | NA |
| S2-7$_{dimer}$ | S2 | 246 |
| S2-10 | S2 | 1718 |
| S2-10$_{dimer}$ | S2 | 20.4 |

NA - No activity
