## Supplementary Table 4 for "Nanobody Repertoires for Exposing Vulnerabilities of SARS-CoV-2"

| ID | Spike S1 variant | $K_{on}$ ($M^{-1}s^{-1}$) | $K_{off}$ ($s^{-1}$) | $K_D$ (M) |
|---|---|---|---|---|
| S1-1 | WT (Wuhan 2019) | 4.14E+05 | 2.98E-05 | 7.20E-11 |
| | 20I/501Y.V1 | 2.71E+05 | 1.21E-05 | 4.44E-11 |
| | 20H/501Y.V2 | 2.78E+05 | 1.25E-05 | 4.51E-11 |
| S1-6 | WT (Wuhan 2019) | 1.02E+06 | 5.75E-04 | 5.65E-10 |
| | 20I/501Y.V1 | 3.55E+06 | 6.27E-04 | 1.77E-10 |
| | 20H/501Y.V2 | 1.03E+06 | 3.29E-04 | 3.20E-10 |
| S1-23 | WT (Wuhan 2019) | 2.82E+06 | 4.91E-05 | 1.74E-11 |
| | 20I/501Y.V1 | 5.96E+06 | 1.36E-03 | 2.29E-10 |
| | 20H/501Y.V2 | NA | NA | NA |
| S1-RBD-9 | WT (Wuhan 2019) | 2.85E+05 | 1.23E-04 | 4.30E-10 |
| | 20I/501Y.V1 | 4.84E+04 | 3.88E-05 | 8.01E-10 |
| | 20H/501Y.V2 | 1.34E+05 | 9.55E-05 | 7.13E-10 |
| S1-RBD-11 | WT (Wuhan 2019) | 2.22E+07 | 2.94E-04 | 1.32E-11 |
| | 20I/501Y.V1 | 3.84E+06 | 2.87E-04 | 7.46E-11 |
| | 20H/501Y.V2 | 6.85E+06 | 1.10E-03 | 1.61E-10 |
| S1-RBD-15 | WT (Wuhan 2019) | 5.37E+06 | 1.50E-04 | 2.79E-11 |
| | 20I/501Y.V1 | 2.99E+06 | 1.26E-04 | 4.22E-11 |
| | 20H/501Y.V2 | 4.02E+06 | 2.22E-04 | 5.53E-11 |
| S1-RBD-35 | WT (Wuhan 2019) | 8.01E+05 | 1.68E-04 | 2.10E-10 |
| | 20I/501Y.V1 | 1.33E+07 | 2.40E-03 | 1.80E-10 |
| | 20H/501Y.V2 | 5.94E+06 | 2.65E-03 | 4.46E-10 |

NA - No activity
