## Supplementary Table 5 for "Nanobody Repertoires for Exposing Vulnerabilities of SARS-CoV-2"

| ID | Epitope | SARS-CoV-2 PSV IC50 (s.e.m) (nM) | SARS-CoV-1 PSV IC50 (s.e.m) (nM) | SARS-CoV-2 20H/501Y.V2 PSV IC50 (s.e.m.) (nM) |
|---|---|---|---|---|
| S1-1 | RBD | 6.74 (1.03) | 8.62 (6.38) | 5.83 (1.40) |
| S1-3 | non-RBD | 1030 (666) | 3,598 (563) | – |
| S1-4 | RBD | 41.5 (3.68) | 179 | – |
| S1-6 | RBD | 56.1 (20.7) | 227 (205) | – |
| S1-17 | non-RBD | 1271 (888) | NA | – |
| S1-20 | RBD | 51.8 (3.72) | NA | NA |
| S1-23 | RBD | 5.69 (2.16) | NA | NA |
| S1-24 | non-RBD | 868 | NA | – |
| S1-27 | RBD | 19.5 (4.90) | NA | – |
| S1-30 | non-RBD | 330 | NA | – |
| S1-31 | RBD | 78.7 (3.48) | NA | – |
| S1-35 | RBD | 12.5 (0.101) | 386.8 (350) | – |
| S1-36 | RBD | 48.5 (21.1) | NA | – |
| S1-37 | RBD | 7.54 | NA | NA |
| S1-39 | RBD | 111 (4.03) | 3.60 | – |
| S1-41 | non-RBD | 732 | NA | – |
| S1-48 | RBD | 5.82 (0.832) | NA | – |
| S1-49 | non-RBD | 356 (56.9) | NA | – |
| S1-51 | RBD | 608 | NA | – |
| S1-58 | non-RBD | 146 | NA | – |
| S1-62 | RBD | 4.95 | – | NA |
| S1-RBD-6 | RBD | 77.2 (21.8) | 89.7 | – |
| S1-RBD-9 | RBD | 522 | – | 116 |
| S1-RBD-11 | RBD | 13.5 (5.50) | NA | 42.9 |
| S1-RBD-15 | RBD | 5.29 (1.44) | NA | 0.273 |
| S1-RBD-16 | RBD | 79.2 (4.23) | – | 1,612 |
| S1-RBD-20 | RBD | 12.4 (1.06) | NA | NA |
| S1-RBD-21 | RBD | 14.1 | – | 115 |
| S1-RBD-23 | RBD | 7.31 (0.418) | NA | 18.5 |
| S1-RBD-24 | RBD | 218 | 12.3 | – |
| S1-RBD-27 | RBD | 65.8 | NA | NA |
| S1-RBD-29 | RBD | 9.53 (1.04) | NA | – |
| S1-RBD-35 | RBD | 12.3 (2.40) | NA | 51.2 |
| S1-RBD-37 | RBD | 391 | – | NA |
| S1-RBD-40 | RBD | 25.6 (5.88) | – | 500 |
| S1-RBD-47 | RBD | 127 (11.6) | – | 112 |
| S1-RBD-48 | RBD | 54.5 | – | NA |
| S2-3 | S2 | 2,234 (751) | 6,277 | – |

"–" - Not determined

NA - No activity
