## Supplementary Table 6 for "Nanobody Repertoires for Exposing Vulnerabilities of SARS-CoV-2"

| protein1 | peptide1 | residue1 | protein2 | peptide2 | residue2 |
| --- | --- | --- | --- | --- | --- |
| S1-1 | DNAK*NTAYLQMNSLKPEDTAVYYCAADSK | 80 | RBD | QIAPGQTGK*IADYNYK | 417 |
| S1-1 | QAPGK*ER | 47 | RBD | K*SNLKPFER | 458 |
| S1-1 | QAPGK*ER | 47 | RBD | SNLK*PFER | 462 |
| S1-1 | QAPGK*ER | 47 | RBD | QIAPGQTGK*IADYNYK | 417 |
| S1-1 | TASRDNAK*N | 80 | RBD | GVSPTK*LNDLCF | 386 |
| S1-1 | LADSVK*GRF | 69 | RBD | RK*SNLKPF | 458 |
| S1-1 | LADSVK*GRF | 69 | RBD | SK*VGGNY | 444 |
| S1-1 | TASRDNAK*NTAY | 80 | RBD | NSNNLDSK*VGGNY | 444 |
| S1-1 | LQMNSLK*PEDTAVY | 91 | RBD | NSNNLDSK*VGGNY | 444 |
| S1-1 | TASRDNAK*N | 80 | RBD | RK*SNLKPF | 458 |
| S1-1 | TASRDNAK*NTAY | 80 | RBD | VIRGDEVRQIAPGQTGK*IADY | 417 |
| S1-1 | LADSVK*GRF | 69 | RBD | VIRGDEVRQIAPGQTGK*IADY | 417 |
| S1-1 | YCAADSK*GW | 105 | RBD | GVSPTK*L | 386 |
| S1-23 | EDSVK*GRF | 69 | RBD | NSNNLDSK*VGGNY | 444 |
| S1-23 | EDSVK*GRF | 69 | RBD | RKSNLK*PF | 462 |
| S1-23 | QAPGK*ER | 47 | RBD | SNLK*PFER | 462 |
| S1-23 | RQAPGK*EREF | 47 | RBD | NSNNLDSK*VGGNY | 444 |
| S1-23 | QAPGK*ER | 47 | RBD | SNLK*PFER | 462 |
| S1-23 | QMNSLK*PE | 91 | RBD | APGQTGK*IADY | 417 |
| S1-23 | MNSLK*PEDTAVY | 91 | RBD | K*VGGNYNYLYRLF | 444 |
| S1-23 | K*PEDTAVY | 91 | RBD | EVRQIAPGQTGK*IADY | 417 |
| S1-RBD-15 | QAPGK*ER | 47 | RBD | K*SNLKPFER | 458 |
| S1-RBD-15 | QAPGK*ER | 47 | RBD | QIAPGQTGK*IADYNYK | 417 |
| S1-RBD-15 | TISRDNAK*NTVY | 80 | RBD | NSNNLDSK*VGGNY | 444 |
| S1-RBD-15 | TISRDNAK*N | 80 | RBD | RK*SNLKPF | 458 |
| S1-RBD-15 | TISRDNAK*N | 80 | RBD | GVSPTK*LNDLCF | 386 |
| S1-RBD-15 | QAPGK*ER | 47 | RBD | SNLK*PFER | 462 |
| S1-RBD-15 | TISRDNAK*NTVY | 80 | RBD | NRK*RISNCVADY | 356 |

**Supplementary Table 6. DSS-Crosslinked Nanobody-RBD Peptides Used for Modeling.** S1-1, S1-23, and S1-RBD-15 nanobodies were bound to RBD and crosslinked with DSS (disuccinimidyl suberate). Crosslinked complexes were excised from SDS-PAGE gels, reduced, alkylated, and digested with either trypsin or chymotrypsin. Peptides were extracted and analyzed by mass spectrometry. Crosslinked peptides (listed) and residues (indicated by asterisk) were identified using pLink, and spectra were manually validated to eliminate false positives.
