## Supplementary Table 7 for "Nanobody Repertoires for Exposing Vulnerabilities of SARS-CoV-2"

**Nanobody neutralization of rVSV/SARS-CoV2 and selected resistant mutants**

| Nanobody | Epitope | rVSV/SARS-CoV-2 variant | IC50 (nM) ± SEM |
| --- | --- | --- | --- |
| S1-1 | RBD | wt | 2.63 ± 0.23 |
|  |  | **Y369N** | 122 ± 3.0 |
|  |  | **G404E** | 40.8 ± 1.01 |
| S1-6 | RBD | wt | 13.0 ± 3.47 |
|  |  | **D574N**, Q792H, Q992H | 587 ± 31.1 |
|  |  | **S371P**, H66R, N969T | 202 ± 29.9 |
| S1-23 | RBD | wt | 0.58 ± 0.02 |
|  |  | **F490S**, **E484K**, **Q493K** | >1,000 |
|  |  | **Q493R**, G252R | >1,000 |
| S1-36 | RBD | wt | 3.69 ± 0.14 |
|  |  | W64R,**L452F** | 262 ± 10.1 |
|  |  | H245R, H1083Y | 2.65 ± 0.24 |
|  |  | W64R,**F490L**,I931G | 870 ± 202 |
|  |  | W64R, **F490S** | >1,000 |
| S1-37 | RBD | wt | 1.83 ± 0.59 |
|  |  | W64R, **F490S** | >1,000 |
| S1-48 | RBD | wt | 1.75 ± 0.43 |
|  |  | **Y449H**, **F490S**, Q787R | >1,000 |
|  |  | **S494P** | >1,000 |
| S1-49 | S1 non-RBD | wt | 146 ± 53.8 |
|  |  | **S172G** | >1,000 |
| S1-49_dimer_ |  | wt | 3.38 ± 2.44 |
|  |  | **S172G** | >1,000 |
| S1-49_trimer_ |  | wt | 0.47 ± 0.00 |
|  |  | **S172G** | >1,000 |
| S1-62 | RBD | wt | 0.65 ± 0.16 |
|  |  | **E484K** | >1,000 |
| S2-10 | S2 | wt | 6,649 ± 2,545 |
|  |  | W64R, **S982R** | >100,000 |
| S2-10_dimer_ |  | wt | 1015 ± 236 |
|  |  | W64R, **S982R** | >40,000 |
| S1-RBD-9 | RBD | wt | 30.2 ± 7.43 |
|  |  | T259K, **K378Q** | >1,000 |
|  |  | W64R, **K378Q** | >1,000 |
|  |  | **K378Q** | >1,000 |
| S1-RBD-11 | RBD | wt | 1.44 ± 0.53 |
|  |  | **F486S** | >1,000 |
|  |  | **T478R** | >1,000 |
|  |  | **T478I** | >1,000 |
| S1-RBD-15 | RBD | wt | 1.21 ± 0.06 |
|  |  | **Y508H** | 549 ± 36.9 |
| S1-RBD-16 | RBD | wt | 268 ± 162 |
|  |  | **N354S** | >1,000 |
| S1-RBD-21 | RBD | wt | 9.61 ± 1.90 |
|  |  | **F486L** | >1,000 |
|  |  | **Y489H** | >1,000 |
| S1-RBD-22 | RBD | wt | 31.5 ± 11.8 |
|  |  | **K378Q** | >1,000 |
| S1-RBD-23 | RBD | wt | 14.8 ± 3.55 |
|  |  | **L452R** | >1,000 |
|  |  | H245R, **S349P**, H1083Y | >1,000 |
| S1-RBD-24 | RBD | wt | 58.0 ± 0.00 |
|  |  | **P384Q** | >1,000 |
| S1-RBD-29 | RBD | wt | 18.0 ± 1.80 |
|  |  | **E484G** | >1,000 |
|  |  | **E484K** | >1,000 |
| S1-RBD-35 | RBD | wt | 1.80 ± 0.15 |
|  |  | **T478I** | >1,000 |
| S1-RBD-40 | RBD | wt | 38.9 ± 11.7 |
|  |  | **K378Q** | 11.8 ± 17.7 |
|  |  | W64R, **F490S** | >500 |
